# Adaptations in gut Bacteroidales facilitate stable co-existence with their lytic bacteriophages

**DOI:** 10.1101/2024.11.17.624012

**Authors:** Adrián Cortés-Martín, Colin Buttimer, Jessie L. Maier, Ciara A. Tobin, Lorraine A. Draper, R. Paul Ross, Manuel Kleiner, Colin Hill, Andrey N. Shkoporov

## Abstract

**Background:** Bacteriophages (phages) and bacteria within the gut microbiome persist in long-term stable coexistence. These interactions are driven by eco-evolutionary dynamics, where bacteria employ a variety of mechanisms to evade phage infection, while phages rely on counterstrategies to overcome these defences. Among the most abundant phages in the gut are the crAss-like phages that infect members of the Bacteroidales, in particular *Bacteroides*. In this study, we explored some of the mechanisms enabling the co-existence of four phage-Bacteroidales host pairs *in vitro* using a multi-omics approach (transcriptomics, proteomics and metabolomics). These included three *Bacteroides* species paired with three crAss-like phages (*Bacteroides intestinalis* and ϕcrAss001, *Bacteroides xylanisolvens* and ϕcrAss002, and an acapsular mutant of *Bacteroides thetaiotaomicron* with DAC15), and *Parabacteroides distasonis* paired with the siphovirus ϕPDS1.

**Results:** We show that phase variation of individual capsular polysaccharides (CPSs) is the primary mechanism promoting phage co-existence in Bacteroidales, but this is not the only strategy. Alternative resistance mechanisms, while potentially less efficient than CPS phase variation, can be activated to support bacterial survival by regulating gene expression and resulting in metabolic adaptations, particularly in amino acid degradation pathways. These mechanisms, also likely regulated by phase variation, enable bacterial populations to persist in the presence of phages, and *vice versa*. An acapsular variant of *B. thetaiotaomicron* demonstrated broader transcriptomic, proteomic, and metabolomic changes, supporting the involvement of additional resistance mechanisms beyond CPS variation.

**Conclusions:** This study advances our understanding of long-term phage-host interaction, offering insights into the long-term persistence of crAss-like phages and extending these observations to other phages, such as ϕPDS1. Knowledge of the complexities of phage-bacteria interactions is essential for designing effective phage therapies and improving human health through targeted microbiome interventions.

## 1. Introduction

The human gastrointestinal tract (GIT) harbours a complex and diverse population of microorganisms (bacteria, archaea, eukarya) and their viruses, which exhibit a high degree of taxonomic diversity [1]. These inhabitants have been implicated in human health and diseases since they can be directly or indirectly involved in essential physiological and metabolic functions within the GIT [1, 2,3,4]. Bacteriophages (phages), viruses that infect bacteria, are one of the most abundant entities within the gut microbiome [4]. From an ecological standpoint, phages are commonly seen as predators that bacteria actively evade, regardless of the mode of phage infection (lytic, lysogenic, or chronic infection) [5]. Bacteria constantly acquire new resistance mechanisms against viruses to evade infection, while phages evolve strategies to overcome these defensive mechanisms [6,7,8]. This selective pressure constitutes a force influencing ecological and evolutionary dynamics within microbial communities called antagonistic coevolution [9,10,11]. This phenomenon leads to diversity in both phage and bacterial populations in natural environments, resulting in a mutually beneficial outcome that secures the perpetuation of both partners without dramatic fluctuations and extinction events [11]. There are two main biological models used to explain phage-host coevolution in the human gut; the ‘arms race’ dynamic that involves continuous directional selection of mutations leading to expanded ranges of host and phage resistance and infectivity over time, and the fluctuating selection dynamic involving density-dependent fluctuating selection based on a trade-off between benefits of resistance and its metabolic costs [10,12,13]. The latter model appears to be favoured in the gut, as it allows for localised co-evolution promoting specialised bacterial resistance, narrowing phage host range, and increasing bacterial diversity, as is usually described in this environment [14]. However, understanding how this system operates in the human gut to perpetuate phage and bacterial survival and its implications for human health is still an area of ongoing research.

The crAss-like phages (order *Crassvirales*) are one of the most abundant groups of dsDNA-tailed phages in the human gut [15]. Their bacterial hosts belong to the order Bacteroidales, which is one of the most prevalent bacterial taxonomic groups in the healthy human gut microbiota, where *Bacteroides* and *Parabacteroides* are two of the most abundant genera [15,16]. To date, only a limited number of gut crAss-like phages have been successfully isolated in the laboratory [17,18,19,20,21], and even fewer have undergone thorough biological characterisation [17,19]. Despite their virulent nature, crAss-like phages can effectively co-exist with their host bacteria without dramatically impacting community structure or target bacterial numbers [19,22,23]. Human gut Bacteroidales bacteria often contain multiple loci encoding for diverse capsular polysaccharides (CPSs). These surface structures are implicated in several functions, such as evading or modifying the host immune response, protecting bacterial cells against phage, and facilitating bacterial colonization and adhesion [24,25]. This molecular mechanism also supports the concept of fluctuating selection dynamics between phage and host among species of Bacteroidetes. Dynamic phase variation in CPS expression has been proposed as an explanation for the rapid acquisition of phage resistance within a high proportion of the host population, as seen in the case of ϕcrAss001 infecting *B. intestinalis* [22]. However, further in-depth investigations with other crAss-like phages and other phage types infecting Bacteroidales are lacking. Additionally, the co-existence observed between ϕcrAss001 and its host bacterium may not be a unique characteristic of *Crassvirales*. Many tailed phages demonstrate a similar pattern of relatively supportive cohabitation with their hosts over numerous generations in the mammalian gut [15].

Here, we investigated the mechanisms underlying phage-bacteria co-existence in bacteria and phage isolated from the mammalian gut through an *in vitro* multi-omics approach. We investigated four phage-bacteria pairs: three *Bacteroides* strains supporting replication of three distinct crAss-like phages (*Bacteroides intestinalis* APC919/174 and ϕcrAss001, *Bacteroides xylanisolvens* APCS1/XY and ϕcrAss002, *Bacteroides thetaiotaomicron* VPI-5482 *tdk*^-^ engineered acapsular mutant [Δcps] and DAC15), and *Parabacteroides distasonis* APCS2/PD with the siphovirus ϕPDS1. Our findings reveal that while CPS phase variation is the primary method that promotes phage co-existence in the Bacteroidales, it is not the only mechanism. Alternative competing resistance mechanisms, which also involve phase variation but less efficiently than changes in CPS expression, can still exert sufficient quantitative effect to allow bacterial populations to survive in the presence of phages. This suggests that bacteria employ multiple strategies to resist phage predation, ensuring their co-existence in the gut microbiome.

## 2. Material and Methods

### 2.1. Bacteriophages, bacterial hosts, and culture conditions

Four phage-bacteria pairs were investigated in this study. The host strains evaluated were: *Bacteroides intestinalis* APC919/174 (RefSeq accession number of complete genome NZ_CP041379), *Bacteroides xylanisolvens* APCS1/XY (RefSeq: NZ_CP042282), *Parabacteroides distasonis* APCS2/PD (RefSeq: NZ_CP042285), and *Bacteroides thetaiotaomicron* VPI-5482 *tdk^-^*acapsular [Δcps] (RefSeq: NC_004663), and along with their corresponding phages ϕcrAss001 (Refseq: NC_049977), ϕcrAss002 (Refseq: NC_055828), ϕPDS1 (Genbank: MN929097 [RefSeq not available]), and DAC15 (RefSeq: NC_055832), respectively. Frozen pure stocks of the host strains stored in 30% glycerol at −80 °C were streaked on Fastidious Anaerobe Agar (FAA) (Neogen) plates (1.5% agar w/v) and incubated at 37 °C for 48 hours in anaerobic jars (Thermo Fisher Scientific) with AnaeroGen^TM^ Compact (Thermo Fisher Scientific) to obtain isolated colonies to perform each experiment. Overnight cultures were obtained after picking a single bacterial colony grown on FAA plates and inoculating into 10 mL of Fastidious Anaerobe Broth (FAB) (Neogen) and incubating overnight at 37 °C anaerobically in anaerobic jars or in a type A vinyl anaerobic chamber (Coy Laboratory Products).

### 2.2. Phage-bacteria *in vitro* co-culture experiment

Co-cultures of each host with its phage were performed in triplicate by serial sub-culturing in broth over five days. Controls which were not infected were also carried out in triplicate. This was accomplished by picking single bacterial colonies, inoculating them into 10 mL of FAB and incubating them overnight at 37 °C anaerobically. The next day, 200 µL of the overnight culture was inoculated in 10 mL of fresh FAB. When bacteria reached the early log phase (OD_595_ = ∼0.2-0.3), they were infected with a corresponding phage at a multiplicity of infection (MOI) of 1 and cultured for 24 hours at 37 °C in anaerobic conditions. An equal volume of growth media without phage was added to the controls. Subsequent rounds of sub-culturing were conducted daily for five days by introducing the previous day’s co-culture into 10 mL fresh FAB medium at a ratio of 1:50. Bacterial and phage enumeration was carried out daily following the overnight incubation. This involved plating bacterial dilutions and performing plaque assay or qPCR for phage counting. Bacteria were counted by serial diluting of 500 µL of overnight culture in phosphate-buffered saline (PBS) (Sigma-Aldrich). 100 µL of the diluted sample was spread on FAA agar plates (1.5% agar w/v) and incubated in anaerobic conditions at 37 °C for 48 hours. The number of generations per day was calculated using the formula: N = N_0_ x 2^n^, where n is the number of generations, N is the number of cells after 24 hours, and N_0_ is the initial number of cells at the start of the day. For phage counting, the overnight culture was centrifuged to remove the cells at 4,500 x *g* for 20 min at 4 °C. The supernatant was filtered through a 0.45 µm pore syringe-mounted polyethersulfone (PES) membrane filter (Sarstedt). The quantification of ϕcrAss001 and DAC15 was performed through plaque assays by mixing 200 µL of an overnight host bacteria culture, 100 µL of SM buffer (1 M Tris HCl pH 7.5, 5 M NaCl, 1 M MgSO_4_) diluted phage supernatant and 3 mL of 0.45% FAA molten overlay soft agar with the addition of MgSO_4_ and CaCl_2_ (1 mM final concentration) and poured onto FAA agar plates (1.5% agar w/v). Since ϕPDS1 does not produce clear plaques that can be counted with the naked eye, and ϕcrAss002 does not produce individual plaques, both phages were quantified by qPCR with the standard curve method using the same conditions and primers described previously [19,26].

To assess the potential influence of the initial MOI on the infection, a range of different initial MOIs (0.001 – 10) was tested for each phage-host pair, following the same procedure described above for the co-culture experiment over five days.

A growth curve for all phage-host pairs under the tested conditions was performed on the fifth day of the co-culture experiment to evaluate potential variations in bacterial growth in the presence of phage. A 1/50 dilution of the fifth-day co-culture in fresh media was prepared. Aliquots of 200 µL, in triplicate, were dispensed into the wells of a flat-bottom 96-well micro test plate (Sarstedt). The plate was sealed, and the OD_595_ was measured under anaerobic conditions for 24 hours using a microtiter plate reader (Multiskan FC, Thermo Fisher Scientific). FAB was used as a negative control. All samples were analysed in triplicate.

### 2.3. RNA extraction, RNA sequencing, and RNA-seq analysis

Samples for the transcriptomics analysis were collected during the bacterial mid-log phase on the fifth day of the co-culture experiment, which occurred between 5-7 hours after the subculture (OD_595_= ∼0.2-0.3). For each of the phages examined in the study, three control samples (consisting only of bacteria) and three phage-bacteria co-culture samples were analysed. One millilitre of the mid-log phase culture was transferred to RNase-free tubes (Thermo Fisher Scientific) and centrifuged for 3 min at 17,000 x *g*. The pellet was resuspended and lysed in 1 mL of TRIzol™ Reagent (Life Technologies), and the total bacterial RNA was extracted into a final volume of 30 µL following the manufacturer’s guidelines. The RNA samples (30 µL) were treated with the TURBO DNA-free kit (Thermo Fisher Scientific) to remove any DNA contamination and were once again purified and concentrated using the RNeasy Mini kit (Qiagen) following the manufacturer’s protocol. The quantity, purity, and integrity of the isolated RNA were measured by Qubit with the RNA High Sensitivity assay (Thermo Fisher Scientific), Nanodrop (Thermo Fisher Scientific), and Bioanalyzer 2100 (Agilent Technologies) with the Agilent RNA 6000 nano kit (Agilent Technologies), respectively. Strand-specific RNA-Seq libraries were prepared and sequenced by Genewiz (Leipzig, Germany) on the Illumina NovaSeq 6000 platform using a 2 x 150 bp paired-end sequencing configuration. RNA-Seq analysis was conducted following the practice guidelines proposed by Love et al. (2015) [27]. Briefly, RNA-seq reads were mapped to host and phage reference genomes [*Bacteroides intestinalis* APC919/174 (RefSeq: NZ_CP041379) and ϕcrAss001 (Refseq: NC_049977), *Bacteroides xylanisolvens* APCS1/XY (RefSeq: NZ_CP042282) and ϕcrAss002 (Refseq: NC_055828), *Parabacteroides distasonis* APCS2/PD (RefSeq: NZ_CP042285) and ϕPDS1 (Genbank: MN929097), and *Bacteroides thetaiotaomicron* VPI-5482 (RefSeq: NC_004663) and DAC15 (RefSeq: NC_055832)] using Bowtie2 (v2.5.1) in end-to-end alignment mode [28]. SAMtools was used to process the resulting alignment SAM files to obtain sorted BAM files. A read count matrix, summarising transcript abundance on a per-gene basis, was constructed using the *summarizeOverlaps* function from the R package GenomicAlignments. Differential gene expression analysis was carried out on this count matrix using the DESeq2 package in R [27].

### 2.4. Ratio of sensitive-resistant bacterial colonies after phage infection

The dynamics of the sensitive/resistant cell ratios in phage-bacteria co-cultures were examined for the four bacterial strains tested in this study. Early log phase cultures (OD_595_= ∼0.2-0.3) were infected with their respective phages at an MOI of 1 and incubated anaerobically at 37 °C for 24 hours. Negative controls, consisting of bacteria without phage infection, were also included. Daily subcultures (dilution 1:50) in fresh media were performed for five consecutive days. On each day, 200 µL of serially diluted overnight culture was mixed with 3 mL of FAB soft agar (0.45%) and 100 µL of a high-titre phage stock (>10^9^) of the host. The mixture was poured onto FAA (1.5%) plates and incubated under anaerobic conditions at 37 °C for 48 hours. Overnight cultures with no extra phage added were also plated and incubated under the same conditions. The percentage of resistant colonies was determined by dividing the number of colonies counted when additional phage was added (resistant sub-population) by the number of colonies counted on the plates where phage was not added (total population).

### 2.5. *In silico* analysis

Long-read sequencing data of *Bacteroides xylanisolvens* APCS1/XY from Guerin et al. (2021) [19] were used to identify potential structural variants within its genome. To accomplish this, long-read DNA sequences were aligned against the reference genome using minimap2 [29], and putative structural variants were detected using Sniffles (v2.0.7) [30]. The isolation of reads that assembled genomic structural variants were extracted using Samtools (v1.18) [31] and QIIME2 (v2023.7) using the filter_fasta.py script [32]. The identification of reads that supported alternative genomic structures was manually reviewed using Artemis (v16) at genomic coordinates, as indicated by Sniffles analysis, which identified the specific region sequences involved in the structural variants. The relevant detected sequence was then searched within all genomes using Artemis and BLASTn to check if the sequence was a common element within the genome of interest.

A Clusters of Orthologous Groups (COGs) analysis was carried out to classify and categorise protein functions of genes significantly different in the RNA-seq analysis to identify patterns and trends in protein function [33]. Determination of the function of bacterial orthologous groups was performed by RPS-BLAST (v2.14.1) [34] against the COG database (22 January 2018).

The representation of the gene clusters was performed using the clinker tool [35].

### 2.6. Proteomics analysis

Samples for the proteomics analysis were collected during the bacterial mid-log phase (OD_595_= ∼0.2-0.3) on the fifth day of co-culture. For each of the phages examined in the study, four control samples (consisting only of bacteria) and four phage-bacteria co-culture samples were analysed. Bacterial cultures were centrifuged at 4,500 x *g* for 15 min at 4 °C. The supernatant and pellet were stored separately at −80 °C, with any remaining liquid removed from the pellet before storage, prior to protein extraction.

Tryptic digests were prepared from all pellet and supernatant samples following a modified version of the manufacturer’s (Protifi) protocol for S-Trap mini devices, in which dithiothreitol (DTT) and iodoacetamide (IAA) were used as the reducing and alkylating agents, respectively. The supernatants and pellets were prepared for digestion on the S-Trap devices using slightly different methods to account for the unique characteristics of each sample matrix. The supernatants (5-8 mL) were first lyophilised to reduce sample volume followed by solubilisation in 1x lysis buffer (5% sodium dodecyl sulfate [SDS], 50 mM triethylammonium bicarbonate [TEAB] pH 7.5) at a ∼1:10 sample to lysis buffer ratio. The pellets contained suspended agar from the growth media which co-pelleted with the bacterial cells. To aid with agar dissolution, 2x lysis buffer (10% SDS, 100 mM TEAB pH 7.5) was used [36] to cover each of the pellets (∼500 µL) prior to pellet resuspension with vortexing. The supernatant and pellet samples were heated to 95 °C for 10 min for lysis followed by a 10 min centrifugation at 21,000 x *g* to remove debris. Samples were reduced with 500 mM DTT (final concentration 22 mM) and incubated at 95 °C for 10 min. Samples were alkylated with 500 mM IAA (final concentration 40 mM) and incubated for 30 min in the dark. The remainder of the digestion followed the manufacturer’s protocol. Protein digestion was performed with 0.8 µg of MS grade trypsin (Thermo Scientific Pierce) and incubation overnight at 37 °C in a wet chamber. The eluted peptides were concentrated in a vacuum centrifuge and acidified with 10% formic acid (FA) for a final percentage of 1% FA. Acidification precipitated and removed residual agar. Acidified samples were filtered with a 10 kDa MWCO PES membrane centrifugal filter. Peptide concentrations were determined with a Pierce Micro BCA assay (Thermo Scientific Pierce) following the manufacturer’s instructions.

Samples were analysed by 1D-LC-MS/MS in two separate runs, one for all supernatant samples and one for all pellet samples. Samples were run in a block-randomised design as previously described [37] with each block containing a full replicate of all species and conditions and two washes between blocks. For each sample, 1,000 ng of peptide was loaded onto a 5 mm, 30 µm ID C18 Acclaim® PepMap100 pre-column (Thermo Fisher Scientific) using an UltiMateTM 3000 RSLCnano Liquid Chromatograph (Thermo Fisher Scientific) for desalting. The pre-column was switched in line with a 75 cm x 75 µm analytical EASY-Spray column packed with PepMap RSLC C18, 2 µm material (Thermo Fisher Scientific) which was heated to 60 °C. The analytical column was connected via an Easy-Spray source to an Exploris 480 hybrid quadrupole-Orbitrap mass spectrometer (Thermo Fisher Scientific). Peptides were separated on the analytical column using a 140 min gradient as previously described [38] and ionised via electrospray ionisation (ESI). MS^1^ spectra were obtained at a resolution of 60,000 on a 380 to 1,600 *m/z* window and fragmented with a normalised collision energy of 27%. MS^2^ spectra were obtained for the 15 most abundant ions in the MS^1^ spectra using a maximum injection time of 50 ms, dynamic exclusion of 25 s, and an exclusion of ions of +1 charge state. Roughly 100,000 MS/MS spectra were acquired per sample.

For protein identification, individual protein sequence databases were created for each phage-host pair. Each database contained all the protein sequences from the bacterial host and phage genome in addition to proteins from the cRAP protein sequence database (http://www.thegpm.org/crap/) containing protein sequences of common laboratory contaminants. The accession numbers for the proteomes in each database are as follows: *B. intestinalis* APC919/174 (GenBank: CP041379.1) and ϕcrAss001 (UniProt: UP000262320), *B. xylanisolvens* APCS1/XY (GenBank assembly: GCA_018279805.1) and ϕcrAss002 (UniProt: UP000595483), *P. distasonis* APCS2/PD (GenBank: CP042285.1) and ϕPDS1 (UniProt: UP000595413), *B. thetaiotaomicron* VPI-5482 (RefSeq: NC_004663.1) and DAC15 (RefSeq: NC_055832.1). For protein identification, MS/MS spectra from each sample were searched against the appropriate database using the Sequest HT and Percolator nodes in Proteome Discoverer version 2.3.0.523 (Thermo Fisher Scientific) as previously described [39]. Only proteins identified with medium or high confidence were retained resulting in an overall False Discovery Rate (FDR) of < 5%. Proteins were quantified by calculating normalised spectral abundance factors (NSAF%) as previously described [39].

### 2.7. Untargeted metabolomics analysis

Untargeted metabolomics was performed on bacterial pellets collected during the bacterial mid-log phase on the fifth day of the co-culture experiment. After reaching OD_595_= ∼0.2-0.3, bacterial cultures were centrifuged at 4,500 x *g* for 15 min at 4 °C. The supernatants were discarded, and the bacterial pellets were frozen in liquid nitrogen and stored at −80 °C until further analysis. The extraction of metabolites and analysis were carried out by MS-Omics (Denmark) following an analytical methodology for the detection of the semi-polar metabolite profile, as follows: 350 μL of ice-cold MilliQ water (∼0 °C), 350 μL of pre-cooled methanol (−20 °C), and 350 μL of pre-cooled chloroform (−20 °C) were added into cell pellet tubes. Sample tubes were vortexed for 1 min, snap-frozen in liquid nitrogen for 1 min, and thawed on ice. The freeze-thaw cycle was repeated two times to induce cell lysis. The samples were centrifuged at 4 °C at 15,000 x *g* for 10 min, and the supernatant (500 μL) was transferred to a new tube. Extracts were evaporated under a gentle stream of nitrogen and reconstituted in 120 μL of mobile phase eluent A (10 mM ammonium formate, 0.1% FA in water) with 10% content B (10 mM ammonium formate, 0.1% FA in MeOH). Reconstituted samples were diluted 1:20 and were analysed using a Thermo Scientific Vanquish LC coupled to a Thermo Q Exactive HF MS. An ESI interface was used as ionisation source. Analysis was performed in positive and negative ionisation mode. The Ultra Performance Liquid Chromatography was performed using a slightly modified version of the protocol described by Doneanu et al. (2011) [40]. Peak areas were extracted using Compound Discoverer 3.3 (Thermo Fisher Scientific). Identification of compounds was performed at four levels: i) Level 1: identification by retention times (compared against in-house authentic standards), accurate mass (with an accepted deviation of 3 ppm), and MS/MS spectra, ii) Level 2a: identification by retention times (compared against in-house authentic standards), accurate mass (with an accepted deviation of 3 ppm), iii) Level 2b: identification by accurate mass (with an accepted deviation of 3 ppm), and MS/MS spectra, and iv) Level 3: identification by accurate mass alone (with an accepted deviation of 3 ppm). To ensure high-quality sample preparation, a quality control sample was prepared by pooling small equal aliquots from each sample to create a representative average of the entire set. This sample was treated and analysed at regular intervals throughout the sequence.

### 2.8. Statistical analysis

Statistical analyses were carried out using the GraphPad Prism (v8.0.1) and R (v2023.12.1+402) software. Data are expressed as the mean ± standard deviation (SD). Normality of data distribution was assessed by the Shapiro-Wilk test. Differences between infected and control samples in experiments (e.g., bacterial counts, % resistant *vs*. sensitive colonies) were evaluated using unpaired t-tests with the Holm-Sidak method for multiple comparisons correction (*p* < 0.05). A two-way ANOVA was applied to analyse the main effects of infection condition, time and their interaction to evaluate growth kinetics, followed by Sidak’s multiple comparisons test for *post-hoc* analysis (*p* < 0.05). For RNA-seq data, differential expression analysis was performed using DESeq2, where generalized linear models were fitted for each gene. The Wald test was used to evaluate significance, and *p*-values were adjusted for multiple comparisons using the Benjamini-Hochberg correction with an FDR set at 5%. The expression values for each CPS cluster were calculated by averaging the expression levels of the genes within each cluster and then normalizing them to the total expression of all CPS clusters. Differences in the relative expression of the CPS loci were analysed assuming normal distribution using Welch’s t-test. In the proteomics analysis, normalised spectral abundance factor (NSAF) values were calculated and log_2_-transformed. Proteins were filtered to include only those with at least three non-zero values across all replicates in both conditions, and missing values were imputed using a normal distribution (width = 0.1, downshift = 2.5). Welch’s t-test was performed followed by Benjamini-Hochberg correction with an FDR set between 5% and 20%. Heat map was generated using z-scored data and Euclidean row and column clustering in R. Volcano plot and principal component analyses (PCA) were also performed using R. For metabolomics, t-tests corrected by the Benjamini-Hochberg method with an FDR of 5% were applied. Multivariate analysis, including PCA and partial least squares-discriminant analysis (PLS-DA), were carried out on metabolites with high accuracy (level 1 and 2a) using the MetaboAnalyst (v6.0) tool [41].

## 3. Results

### 3.1. Stable co-existence of crAss-like phages and ϕPDS1 with their hosts *in vitro*

Four phage-bacteria pairs were co-cultivated by subculturing for five days to investigate the effect of phage on survival and transcriptional responses in *Bacteroides/Parabacteroides* strains over time. Enumeration of bacteria and phages was carried out daily, and it was confirmed that all four phages continued to co-exist with their bacterial hosts over the five days of co-culture, without a complete collapse of either the host or the phage populations (**Fig. 1 A, B, C, D**). The number of generations per day was estimated for each bacterium as ∼6 generations over 24 hours in each experiment, representing 30 generations in total over five days. The dynamics of phage ϕcrAss001 and ϕPDS1 infecting *Bacteroides intestinalis* APC919/174 and *Parabacteroides distasonis* APCS2/PD was similar in that their titre increased after 24 hours of infection, reaching ∼5 × 10^9^ pfu/mL and ∼3 × 10^10^ copies/mL, respectively. In the following days, the titre decreased and stabilized at ∼4 × 10^7^ pfu/mL and ∼5 × 10^8^ copies/mL, respectively (**Fig. 1A, C**). In the case of DAC15 infecting an engineered acapsular mutant of *B. thetaiotaomicron* VPI-5482 (*B. thetaiotaomicron* Δcps), an increase in the titre was observed in the early days, reaching its peak at 48- and 72-hours post-infection with a titre around ∼3 × 10^9^ pfu/mL, which then remained stable beyond the 5-day mark with a titre of ∼2 × 10^8^ pfu/mL (**Fig. 1D**). The absence of CPS phase variation in this strain, previously shown to be the major mechanism of phage evasion [18,24], neither prevented the crAss-like phage DAC15 from propagating at high levels nor did it result in the collapse of bacterial culture, supporting previous data that alternative cell-surface related mechanisms, limiting phage infection through phase variation or otherwise, may be involved [24].

**Fig. 1.**
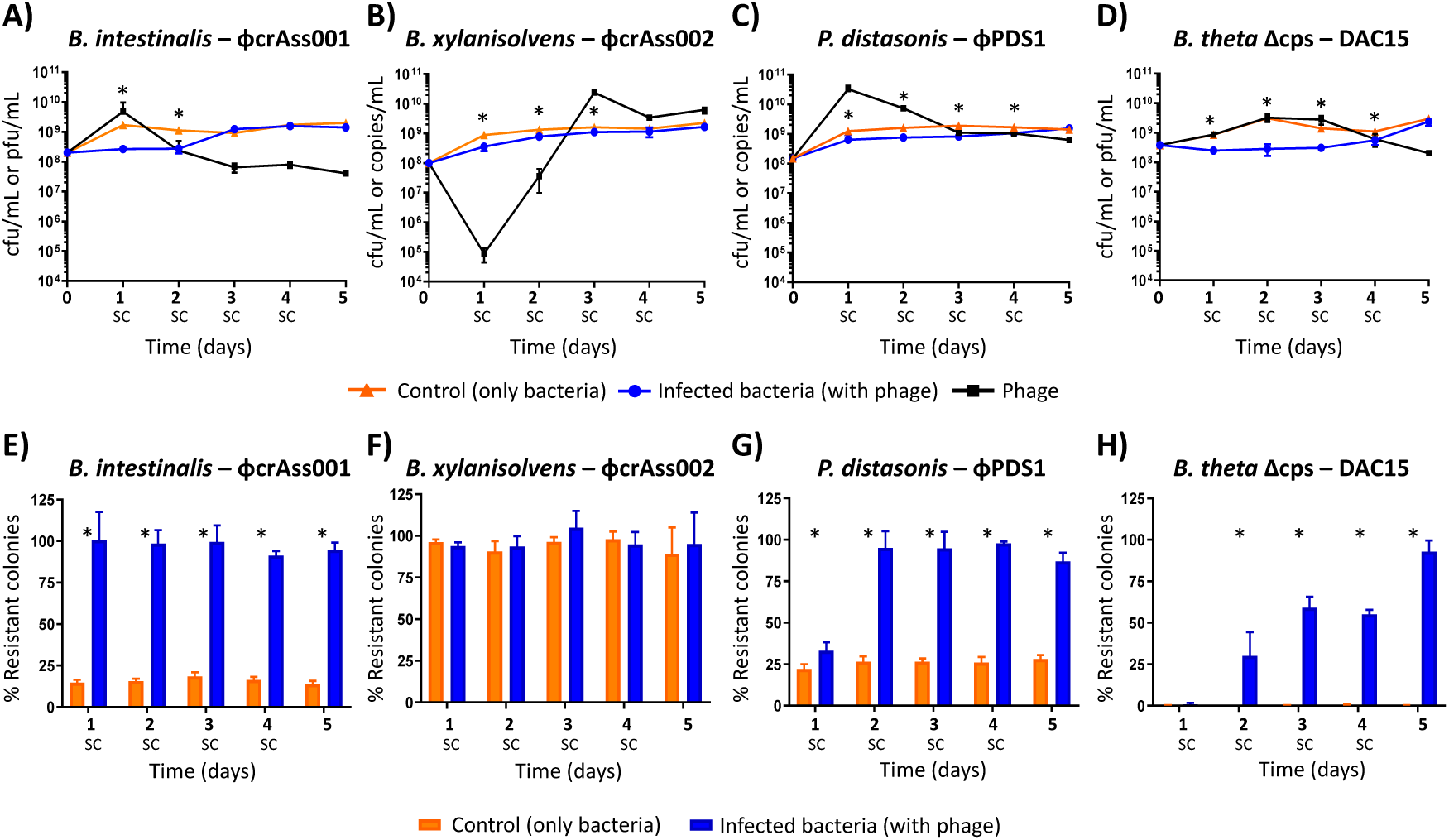
Bacterial-phage dynamics and variation in bacterial resistant colonies acquired during the 5-day co-culture. **(A-D)** Enumeration of bacteria and phage during the 5-day co-culture experiments. After each 24-hour period, a new subculture (SC) was performed in fresh media at a dilution of 1:50. **(A)** *B. intestinalis* + ϕcrAss001, **(B)** *B. xylanisolvens* + ϕcrAss002, **(C)** *P. distasonis* + ϕPDS1, **(D)** *B. thetaiotaomicron* Δcps + DAC15. The orange line represents the bacterial counts (CFU/mL) for the control condition (uninfected bacteria), the blue line represents the bacterial counts (CFU/mL) in co-culture with phage, and the black line represents the phage counts (pfu/mL or copies/mL) in co-culture with bacteria. Differences in phage enumeration methods were due to the ability to produce countable plaques (CFU/mL), or qPCR (copies/mL) when plaques were not detected. **(E-H)** Variation in the percentage of resistant colonies every 24 hours during the 5 days of co-culture experiments. **(E)** *B. intestinalis* + ϕcrAss001, **(F)** *B. xylanisolvens* + ϕcrAss002, **(G)** *P. distasonis* + ϕPDS1, **(H)** *B. thetaiotaomicron* Δcps + DAC15. Orange bars represent the mean percentage of resistant colonies in the uninfected culture (control), while blue bars show the mean percentage of resistant colonies when bacteria were exposed to phage. Error bars indicate standard deviation (SD). All experiments were carried out in triplicate. *: Statistically significant differences in bacterial counts or percentage of resistant colonies when comparing the control with the infected conditions were set at *p* < 0.05.

On the other hand, ϕcrAss002 exhibited very different dynamics. Similarly to a previous report [19], the ϕcrAss002 titre underwent a significant reduction after the initial 24 hours of infection, falling to ∼10^5^ genome copies/mL. However, in the following days, the ϕcrAss002 titre increased considerably, reaching its peak at 72 hours with ∼2 × 10^10^ genome copies/mL and remaining close to this level in the subsequent days (**Fig. 1B**).

Phage co-existence with their hosts was confirmed for all four phages and a titre stabilization around the 72-hour post-infection was observed in all cases except for DAC15, where a slight decrease occurred at the end of the experiment. All phages had a significant impact (*p* < 0.05) on the population of their bacterial hosts in the first days of the experiment, resulting in a reduction in the number of bacteria compared to the control (no infection). These differences diminished during the following days, and by the fifth day, the bacterial values were identical to the control. In *B. intestinalis* and *B. thetaiotaomicron* Δcps, a ∼1 log reduction in bacterial colony-forming unit (CFU) counts was observed between 24-72 hours post-infection compared to the control (**Fig. 1A, D**), which suggests that initially dominant phage-sensitive subpopulations were depleted during the first several rounds of phage infection and were later replaced by less sensitive subpopulations. For *B. xylanisolvens* and *P. distasonis*, the reduction in bacterial counts after initial infection was less pronounced (**Fig. 1B, C**).

The effect of the initial MOI at the moment of the infection was also tested (**Fig. S1**). Over a broad range of MOIs tested (0.001 – 10), no significant differences were observed in the titre of the phages and bacteria at equilibrium after five days of co-culture, despite an MOI-dependent difference in the time taken for the phage-host system to reach an equilibrium (**Fig. S1**). The resulting titres, phage-host ratios at equilibrium, and the ability of phage-host pairs to co-exist stably in co-culture were, therefore, dictated by the properties of phage-host pairs and not by the initial MOI.

The kinetics of bacterial growth were also evaluated after the fifth day of co-culture to determine if the presence of the phage had any impact when phage and bacteria ratios had stabilized. Different kinetics were observed for each phage-host pair (**Fig. S2**). While ϕcrAss001 and DAC15 caused a delay and reduction in the density of the lag phase of their hosts (**Fig. S2A, D**), ϕcrAss002 seemed to have a slight stimulatory effect on the growth of *B. xylanisolvens* (**Fig. S2B**). No significant differences were observed in the kinetics of the growth curve of *P. distasonis* after ϕPDS1 infection (**Fig. S2C**).

### 3.2. Phage exposure increases the percentage of resistant host subpopulations

Phase variation or similar mechanisms leading to phenotypic differentiation and dynamic equilibrium between sensitive and resistant subpopulations [22,24] could explain stable phage-host co-existence in the chosen phage-host pairs. Such dynamics could explain the observed variations in the titre of the phages and their hosts during the five-day experiment. To test this hypothesis, the rate of resistant cells was measured at all time points (**Fig. 1E, F, G, H**). An average of ∼16% of clones in a naïve culture of *B. intestinalis* were found to be resistant to ϕcrAss001 (**Fig. 1E**). However, this number increased to ∼97% after only 24 hours of contact with ϕcrAss001, and a high percentage of resistant clones was then maintained for the remaining days (**Fig. 1E**). This correlates with the rapid drop of phage titre seen in the *B. intestinalis* – ϕcrAss001 co-culture after day 1. In the case of *P. distasonis*, the initial proportion of resistant clones was ∼26% (**Fig. 1G**), rising to ∼35% after 24 hours of ϕPDS1 exposure and then reaching ∼95% after 48 hours, which remained similar until the end of the experiment (**Fig. 1G**). The drop in phage titre was similar to ϕcrAss001, albeit stabilizing at a higher level (10^9^ genome copies/mL compared to 10^7^ pfu/mL) at the 72-hour timepoint.

Unlike the previous two examples, *B. thetaiotaomicron* Δcps showed very different dynamics. The initial ratio of clones resistant to DAC15 was ∼0.3%, indicating high susceptibility to the phage. This value gradually changed to ∼30% at 48 hours, reaching ∼93% on the fifth day (**Fig. 1H**). This result suggests that the lack of the phase-variable CPSs operons necessitates the use by *B. thetaiotaomicron* Δcps of alternative phage defence mechanisms, potentially including phase variation of other surface structures [24], that require more time to build a subpopulation of cells resistant against phage infection. Curiously, the *B. thetaiotaomicron* Δcps-DAC15 pair showed the highest difference in bacterial counts compared to the control (no infection) on days 1-4 during the co-culture experiment, with this resistance appearing to level out on day 5, corresponding to the high level of phage resistance (∼93%) achieved by this time (**Fig. 1D, H**). Such delayed resistance could explain how phage DAC15 maintained high titres during the first three days of co-culturing.

On the other hand, the level of resistance in *B. xylanisolvens* was very high (∼94%) even before contact with phage ϕcrAss002, which potentially explains the inability of ϕcrAss002 to produce visible plaques on this host bacterium [19] (**Fig. 1F**). After exposure to ϕcrAss002, the resistant percentage showed no significant changes over the five days (**Fig. 1F**). This initial high resistance to ϕcrAss002 could explain the differing behaviour of ϕcrAss002 in the *in vitro* co-culture experiment compared to the other phages (**Fig. 1A, B, C, D**). For the other phages a higher percentage of the naïve host cells were sensitive, allowing those phages to propagate and increase in titre during the initial stages of this experiment (**Fig. 1A, B, C, D**).

### 3.3. Phage infection drives dynamic shifts in bacterial surface gene expression through phase variation

After confirming the ability of *Bacteroides* and *Parabacteroides* phages to co-exist with their bacterial hosts without exerting a significant penalty on host cell density, we next examined the transcriptional response of each host to persistent phage presence. RNA-seq analysis was performed for the four phage-bacteria pairs during the mid-log phase of growth on the fifth day of the co-culture experiment.

The transcriptional response for the *B. intestinalis*-ϕcrAss001 pair showed statistically significant changes (*p* < 0.05) in 136 genes (28 downregulated and 108 upregulated), showing mean log_2_ fold changes ranging between −6.14 ± 0.31 and 2.35 ± 0.25 (**Table S1A**). Most of the genes belonged to previously reported phase-variable CPS biosynthesis clusters (operons) termed PVR7, 8, 9, 11 and 12, of which four were upregulated and one (PVR9) – downregulated [22] (**Fig. 2A**). The details of these gene clusters are provided in **Fig. S3A and Table S2A**. These gene clusters shared many common elements. Among them are UpxY family transcription anti-terminators and UpxZ-family inhibitors of UpxY, preventing non-cognate UpxY anti-termination, therefore allowing transcription of only one CPS operon per cell at a time [42,43] Glycosyltransferases and other enzymes are also encoded by these operons that can contribute to CPS chain modifications, such as acetyltransferases and aminotransferases, among others. The transcription of these operons is likely driven by a putative invertible promoter region of 183-186 bp flanked by a conserved 10 bp inverted repeat (IR) GTTCGTTTAA [22]. This strongly suggests that these regions, where recombination is likely to occur, may be relevant in allowing permissive phage infection of the host. As suggested previously, the downregulation of the PVR9 operon as a result of crAss001 exposure is likely to limit opportunities for infection that rely on PVR9 as a phage receptor [22]. At the same time, upregulated CPS operons (7, 8, 11 and 12) may have a protective or neutral effect. Constant switching between expression of different CPS types, therefore, results in a dynamic equilibrium between resistant and sensitive sub-populations, allowing the phage to persist in this mixed population. Clusters of Orthologous Groups (COGs) analysis was performed to classify and categorise the protein functions of the other genes whose expression changed and were located outside the PVR loci, revealing that most of the genes (40%) with known functions were related to inorganic ion transport and metabolism (**Fig. S4A, Table S3A**).

**Fig. 2.**
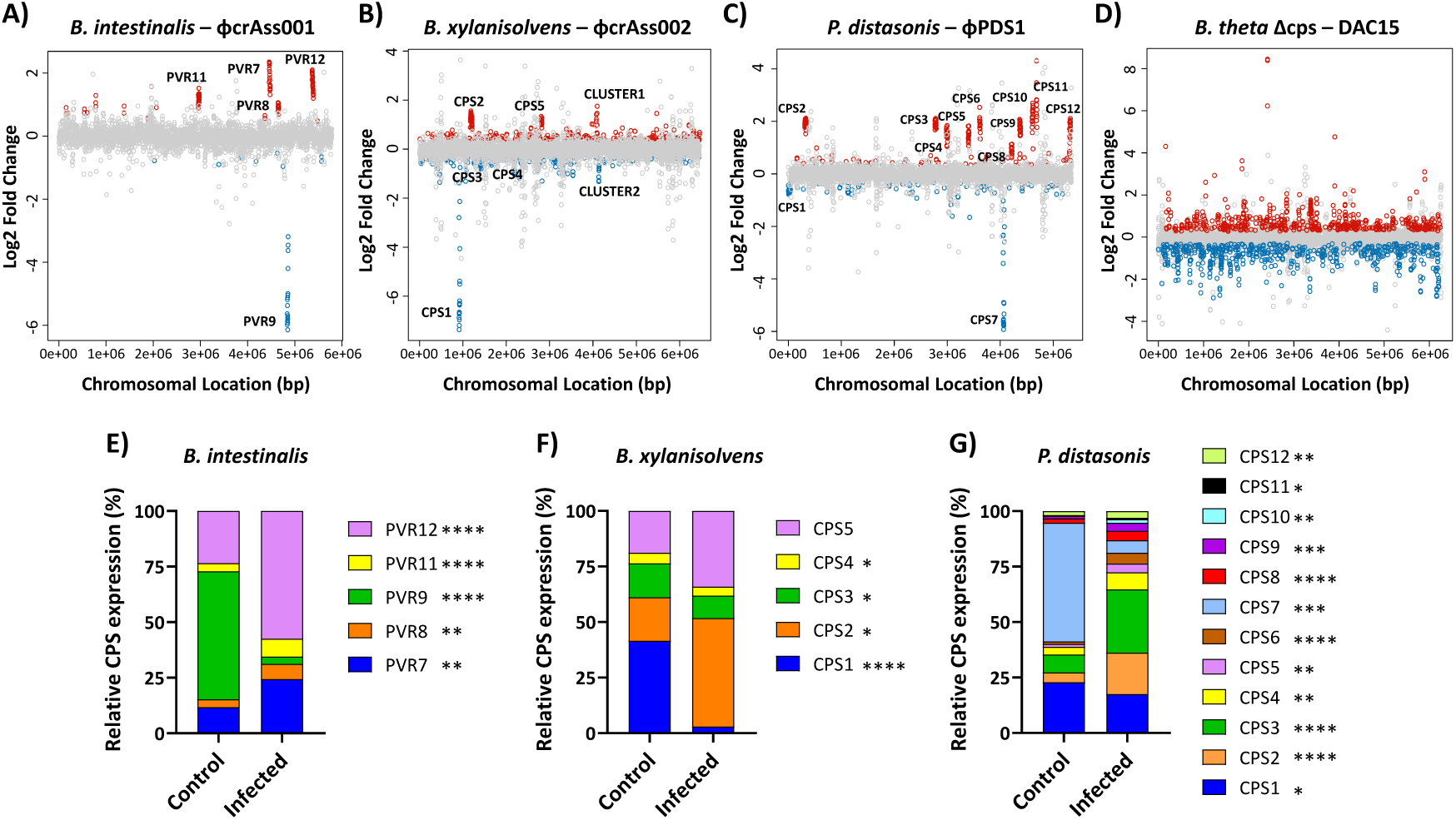
Transcriptional changes and CPS expression profiles after bacteria-phage co-culture. **(A-D)** Transcriptional changes during mid-log phase growth on the fifth day of each bacteria-phage co-culture: **(A)** *B. intestinalis* + ϕcrAss001, **(B)** *B. xylanisolvens* + ϕcrAss002, **(C)** *P. distasonis* + ϕPDS1, **(D)** *B. thetaiotaomicron* Δcps + DAC15. Genes are plotted as dots according to their chromosomal location in each bacterial host. The log_2_ fold change represents the differential expressions of genes when comparing control (uninfected) with infected bacteria. Red dots represent genes that were significantly upregulated upon phage infection, blue dots are those that were significantly downregulated, and grey dots are those that did not show statistically significant changes in expression. Gene clusters implicated in bacterial capsulation are labelled as CPS (capsular polysaccharide) and PVR (phase-variable region). Statistical significance was determined using an FDR-adjusted *p*-value (FDR < 0.05). **(E-G)** Relative expression of the different CPS loci in bacterial capsular strains under infected and uninfected conditions determined by RNA-seq data. **(E)** *B. intestinalis* + ϕcrAss001, **(F)** *B. xylanisolvens* + ϕcrAss002, **(G)** *P. distasonis* + ϕPDS1. The expression values for each CPS cluster were calculated by averaging the expression levels of the genes within each cluster and then normalizing them to the total expression of all CPS clusters. The resulting values are presented as a percentage of the total CPS expression for each condition. Differences in the relative expression of the CPS loci were analysed assuming normal distribution using Welch’s t-test. Significant differences in proportions after phage infection are indicated in the legend as follows: * *p* < 0.05; ** *p* < 0.01; *** *p* < 0.001; **** *p* < 0.0001.

*B. xylanisolvens* showed differences in gene expression of 603 genes (233 downregulated and 370 upregulated) with a mean log_2_ fold change ranging from −7.36 ± 0.14 to 1.75 ± 0.17 (**Table S1B**). Six gene clusters were well differentiated (3 downregulated and 3 upregulated), with the strongest transcriptional response associated with the first identified gene cluster (**Fig. 2B**). Similar to our observations in *B. intestinalis*, four of these gene clusters primarily included genes related to CPS, encoding the UpxY family anti-termination system, glycosyltransferase enzyme genes and other genes related to the biosynthesis of polysaccharides (**Fig. S3B, Table S2B**). Regarding the last two gene clusters, one (CLUSTER1) may represent a Polysaccharide Utilization Loci (PUL), which included components such as the Sus-TonB dependent transporter system, hydrolases, phosphodiesterases, and a DeoR transcription regulator [44], while the other (CLUSTER2) contained genes encoding an ABC transporter, fimbrilin and a TonB-dependent receptor (**Fig. S3B, Table S2B**). Furthermore, variations in gene expression of repetitive genes distributed throughout the genome and not associated with these gene clusters were observed, encompassing genes such as transposases, which may have the potential to induce phase variation, Sus/Rag family and TonB-dependent transporters systems, and ATP-binding proteins (**Table S1B**), which encode bacterial surface molecules that have been previously reported to be implicated in phage sensitivity in *Bacteroides* species [16,22]. The COG analysis for the known functions of genes outside the CPS operons showed more diversity than *B. intestinalis*, with carbohydrate transport and metabolism (13.3%) and amino acid transport and metabolism (10%) being the most relevant categories (**Fig. S4B, Table S3B**). We performed an *in-silico* analysis to detect potential structural variants in this host strain using the long-read genome sequencing data from a previous study [19]. We identified 46 potential structural variants across the *B. xylanisolvens* genome (**Table S4**). However, based on genomic location, only one variant (an inversion) passed all the filters and corresponded to one of the CPS gene clusters detected in our RNA-seq analysis, CPS1. The alignment of the long-read sequences with the reference genome revealed an inversion in the intergenic region of the first gene of the CPS1 operon (FQN58_03730 – FQN58_03810) (**Fig. S5**). Upon further investigation of this operon (**Fig. S5A**), an integrase was identified upstream of the first gene, suggesting potential involvement in genomic recombination within this gene cluster (**Fig. S5B**). A 20 bp IR sequence, GTTACTTCTTAGGTAACGGA, flanked a 290 bp invertible region in the intergenic region preceding the first gene of the CPS1 operon (**Fig. S5C**). This identical repeat sequence was also identified in four other genomic regions, each with an invertible region of size of either 289 bp (in three) or 290 bp (in one). Additionally, an IR exhibiting 90% sequence identity (18/20 bp) (GTTACTTTGTAGGTAACGGA) was identified in both forward and reverse orientations, flanking another potential invertible fragment of a size of 280 bp. Among these matching IRs, three were in the intergenic regions next to the UpxY genes of CPS3 (FQN58_RS05950 – FQN58_RS06080), CPS4 (FQN58_RS09155 – FQN58_RS09165), and CPS5 (FQN58_RS11900 – FQN58_RS11985), and were preceded by a tyrosine-type DNA invertase. The fourth matching fragment was in the intergenic region next to a site-specific integrase (FQN58_RS05740), although no CPS clusters were found nearby. The 18 bp IR was not associated with any of the CPS clusters detected in the RNA-seq analysis but was also preceded by an integrase. The case of the potential CPS4 is noteworthy since it appears to consist of only three genes: an UpxY, a polysaccharide biosynthesis/export family protein and an N-acetylmuramoyl-L-alanine amidase. In the gene coordinate plot with RNA-seq data, this cluster could not be easily distinguished due to the small number of genes involved and a relatively small transcriptional change (**Fig. 2B**). Nevertheless, all three genes within this cluster were found to be downregulated, and the presence of the IR in the intergenic region of the UpxY family antiterminator, preceded by a DNA invertase, suggests that it may represent another phase variable cell surface operon in *B. xylanisolvens*. The remaining CPS operon (CPS2) was preceded by a gene coding for a DUF3078 domain-containing protein, and no repeats or invertases were observed in the intergenic region, unlike in the other CPS operons (**Fig. S3B**). In contrast, this operon includes a transposase as part of its structure. The other two gene clusters identified, which were not associated with CPS structures, were found to be preceded by a transposase (CLUSTER1) and a helix-turn-helix domain-containing protein (CLUSTER2).

The RNA-seq results of the *P. distasonis* – ϕPDS1 pair revealed changes in the expression profile of 393 genes (131 downregulated and 262 upregulated) (**Table S1C**). The mean log_2_ fold change ranged between −5.92 ± 0.36 and 4.30 ± 1.46, and twelve clusters of genes were identified (10 upregulated and 2 downregulated) (**Fig. 2C**). While the two downregulated gene clusters contained the UpxY (CPS1 and CPS7), similar to findings in *B. intestinalis* and *B. xylanisolvens*, the rest of the clusters contained a gene encoding the transcriptional anti-termination protein NusG positioned at the beginning of each operon. This may represent an alternative mode of transcriptional regulation to that of the UpxY anti-terminator system observed in the other *Bacteroides* strains (**Fig. S3C, Table S2C**). The other genes present in these clusters were sugar transferases and modification enzymes similar to the ones described in the other strains, which could also play a role in surface structure changes (**Fig. S3C, Table S2C**). Genes encoding for Sus/Rag family and TonB-dependent transporter systems, polysaccharide biosynthesis proteins and polysaccharide pyruvyl transferase family proteins were other relevant genes whose expression showed significant changes and recurred across the genome, located outside of the mentioned gene clusters (**Table S1C**). COG analysis revealed that most of the genes with a known function outside the CPS operons were categorized under inorganic ion transport and metabolism (20.3%) and cell envelope biogenesis (14.5%) (**Fig. S4C, Table S3C**). We identified that 10 out of 12 CPS operons had a 17 bp consensus IR (GCTACTYRGNRAGTAGC) flanking a putative promoter region of 314 to 345 bp in size. These operons were preceded by a site-specific integrase and corresponded with those containing the anti-termination protein NusG gene in the first position of the operon. Invertible promoter regions flanked by IRs were not found in CPS1 and CPS7. After analysing potential similarities between the IR sequences detected in the three studied strains and those found in other *Bacteroides* species, such as *B. thetaiotaomicron* or *B. fragilis* [24,45], it was observed that, in all cases except for *B. intestinalis*, these IR sequences contained palindromic regions with the same short sequence, GTTAC – GTAAC (with the first thymine replaced by a cytosine in the case of *P. distasonis*), and were also preceded by integrase or invertase enzymes.

These results suggest that alterations in the expression of CPSs are one of the main mechanisms that explain phage-bacteria co-existence in discontinuous culture. The normalized fractions of CPS gene reads mapped to each of the strains changed after phage exposure (*p* < 0.05) (**Fig. 2E, F, G**). The most predominant CPS type in each of the three hosts: PVR9 (57.6%) in *B. intestinalis*, CPS1 (41.5%) in *B. xylanisolvens*, and CPS7 (53.5%) in *P. distasonis* were strongly suppressed to 3.3%, 3.0% and 5.7%, respectively. Simultaneously, the expression of other CPS types was favoured (**Fig. 2E, F, G**), so that PVR12 became the highest expressed CPS (57.4%) in *B. intestinalis*, CPS2 (48.8%) in *B. xylanisolvens*, and CPS3 (28.5%) in *P. distasonis*.

However, CPS gene operons are not the only class of genes involved in bacterial response to phage persistence, as we have seen that DAC15 can also successfully propagate and persist with an acapsular host. The transcriptomics analysis of *B. thetaiotaomicron* Δcps showed the highest number of significantly changed genes among the four phage-bacteria pairs. 1,136 changes were detected, 591 downregulated and 545 upregulated. The mean log_2_ fold change varied from −2.88 ± 0.89 to 8.44 ± 0.57 (**Table S1D**), showing even stronger upregulation of some of the genes compared to the data from capsular strains. Clearly defined gene clusters with a strong transcriptional response could not be identified (**Fig. 2D**). Nevertheless, other groupings of non-consecutive genes were identified. Notably, an upregulation of 30S and 50S ribosomal proteins (BT_RS13645 - BT_RS13840) suggests changes in transcriptional machinery. In addition, other gene groups exhibited changes in expression, including transposases and integrases that may be involved in DNA recombination processes, TonB-dependent transporters and Sus-like systems linked to outer membrane proteins that control the uptake of polysaccharides, and other genes related to membrane proteins like OmpA family proteins or efflux RND transporter permeases, among others (**Table S1D**). Analysis of the COGs revealed a more even distribution among different categories, with no single category prominent. Carbohydrate transport and metabolism was the primary category (15.4%) of known gene functions. In comparison, cell envelope biogenesis and outer membrane category represented 9%, positioning this category as the fourth most relevant in *B. thetaiotaomicron* Δcps (**Fig. S4D, Table S3D**). Most curious was the finding that the most highly upregulated gene, BT_RS09760 (which corresponds to BT1927 in GenBank annotation), is located near a site-specific integrase and has been reported to encode a surface layer (S-layer) protein expressed in a phase-variable manner [46]. Unlike the gene operons of CPSs, which were constituted by multiple genes, only two additional genes (BT_RS09750 - BT_RS09755) were upregulated in this operon. A consensus IR of 19 bp (CCGTTACCTABVRAGTAAC) preceding this operon was also detected in seven other upregulated loci close to site-specific integrases, surface proteins, and OmpA family proteins, as previously reported [24]. Remarkably, a nearly identical palindromic IR sequence (CCGTTACCTAAGAAGTAAC) is found next to tyrosine-type DNA invertases of *B. thetaiotaomicron* wildtype in the CPS operons: CPS1, CPS3, CPS5, and CPS6. This finding shows that these diverse operons are regulated by related mechanisms, enabling phage persistence even in the absence of phase-variable CPS.

This phase variation, regulating CPS and other surface structures, apparently generates frequent and reversible changes within specific hypermutable loci, introducing phenotypic diversity into clonal populations and rendering different populations resistant or sensitive to phage infections [47]. This explains the co-existence of phage and bacteria when they are in co-culture.

### 3.4. Protein-level changes in gene expression in response to phage exposure

Proteomic analysis of bacterial pellets collected during the mid-log phase after five days of daily subcultures with phage exposure did not reveal large differences between phage-infected cultures and negative controls across the bacterial strains tested. Over 2,000 proteins were identified in each strain (**Table S5A, B, C, D**). Principal Component Analysis (PCA) showed no distinct clustering based on experimental conditions (**Fig. S6**), indicating minimal global changes in protein expression due to phage infection. At the individual protein level, only a few proteins showed statistically significant differences with a 5% FDR (**Table S5A, B, C, D**). In *B. intestinalis*, four proteins were differentially abundant: a N-acetyltransferase (WP_007663187.1) belonging to PVR8, an UpxZ family transcription anti-terminator antagonist (WP_115501761.1) and an ATP-grasp domain (WP_115501769.1) both linked to PVR9, and a hypothetical protein (WP_115502230.1) that was not part of the PVR operons (**Table S5A**). *B. xylanisolvens* exhibited differences in the abundance of six proteins: an acyltransferase (WP_055234723.1) associated with CPS2, a polysaccharide export protein (WP_087318117.1) belonging to CPS3, two UpxY family transcription anti-terminators with identical protein sequences (WP_008025447.1) linked to CPS1 and CPS5, and a glycosyltransferase (WP_008643375.1) and a Wzz/FepE/Etk N-terminal domain-containing protein (WP_008643393.1), both belonging to CPS1 (**Table S5B**). *P. distasonis* had only one differentially abundant protein, an electron transport complex subunit D (WP_005855664.1) that was not part of the CPS operons (**Table S5C**), while *Bacteroides thetaiotaomicron* Δcps showed differences in five proteins, members of the PorV/PorQ family (WP_008765856.1), an AAA family ATPase (WP_008766137.1), a BT1926 family outer membrane beta-barrel protein (WP_008766273.1), and two with unknown function (WP_011109276.1 and WP_008762274.1) (**Table S5D**). When applying a less stringent FDR of 20%, additional differentially abundant proteins were identified (**Table S5A, B, C, D**). Comparison with RNA-seq data revealed consistency in some findings. For example, in *B. intestinalis*, 5 of 9 significantly altered proteins coincided with their corresponding genes, also significant in RNA-seq, which belonged to different CPSs: PVR8, PVR9, PVR11. Similarly, in *B. xylanisolvens*, 11 of 12 proteins matched RNA-seq data, all belonging to identified clusters with strong transcriptional responses: CPS1, CPS2, CPS3, and CPS5. No relevant changes were observed in *P. distasonis*, while *B. thetaiotaomicron* Δcps showed 10 out of 19 proteins coinciding with RNA-seq data, including the highly responsive WP_008766274.1 - BT_RS09760 and WP_008766273.1 - BT_RS09755, among others (**Table S5A, B, C, D**).

Supernatant analysis revealed no significant findings for the capsular bacteria (**Fig. S7, Table S6A, B, C**); however, for *B. thetaiotaomicron* Δcps, PCA identified two distinct clusters and 155 differentially expressed proteins at a 5% FDR (**Fig. 3, Table S6D**).

**Fig. 3.**
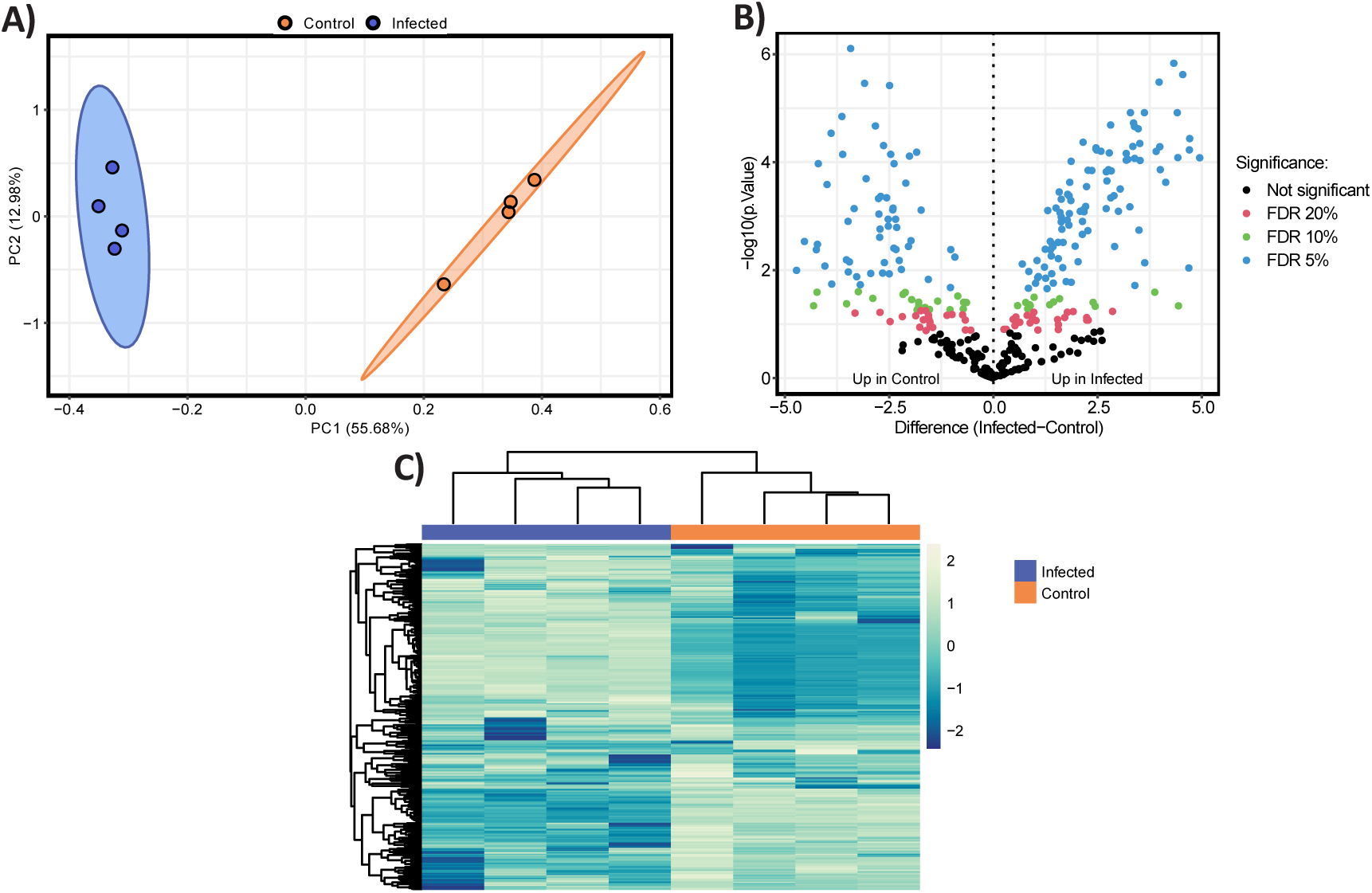
Differential proteomics profile in the supernatant obtained during the mid-log phase on the fifth day of the co-culture experiment with *B. thetaiotaomicron* Δcps and DAC15. **(A)** Principal Component Analysis (PCA) of proteomic profiles of *B. thetaiotaomicron* Δcps supernatant between control (orange) and DAC15-infected samples (blue colour). Ellipses represent 95% confidence intervals, and the explained variances of the two principal components are shown in brackets along the axes. **(B)** Volcano plot representing the significantly differentially abundant proteins. Non-adjusted *p*-values are plotted; however, points are coloured for significance based on corrected *p*-values. Proteins upregulated in control (left panel) and DAC-15 infected (right panel) are plotted based on fold change and statistical significance (Benjamini-Hochberg FDR correction) at different thresholds: 5% (blue), 10% (green), 20% (red), and non-significant (black). **(C)** Heatmap displaying hierarchical clustering of proteomic data generated from z-scored data. Row and columns are clustered based on Euclidean distance, showing differential protein expression between control and infected conditions. Columns are coloured by sample condition with blue representing infected samples and orange representing control. Yellow indicates higher relative protein abundances, while blue represents lower relative abundance.

Surprisingly, 64 of these proteins corresponded to significant changes in the RNA-seq dataset, including the strongly transcriptionally regulated WP_008766274.1 - BT_RS09760, WP_008762274.1 - BT_RS07605 and WP_011109276.1 - BT_RS22595, all of which are associated with the S-layer/OmpA system [24]. Among the proteins not significant in RNA-seq, WP_005677723.1 (encoded by BT_RS00855), a Fe-S protein homologous to NifU, showed the most significant difference in protein abundance. Additionally, 22 of the differentially abundant proteins were ribosomal proteins (30S and 50S), while another 22 were nutrient uptake proteins belonging to the Sus/Rag systems and TonB-dependent transporters, underscoring the relevance of these proteins in mediating DAC15 infection, which also was observed in the RNA-seq analysis. Interestingly, these findings suggest that the acapsular strain exhibits more pronounced proteomic differences compared to the capsular strain, as we also observed in the transcriptomic analysis. While we expected more significant changes in the bacterial pellet, the absence of CPSs may increase interaction with the environment and phages, leading to greater protein release into the supernatant due to lysis.

### 3.5. Intracellular bacterial metabolome reacts to phage exposure with more relevant changes seen in the acapsular strain

To explore whether phage exposure induces metabolic adaptations in the bacterial population that it attacks, we performed untargeted intracellular metabolomics on bacterial pellets during the mid-log phase after five days of co-culture. A total of 1,772 entities were detected by LC-MS, of which 201 were accurately identified with high confidence based on chemical standards (levels 1 and 2a) (**Table S7**). Univariate analysis after 5% FDR correction showed significant changes in the intensity of only two metabolites, 3-methyl-2-oxovaleric acid and 4-methyl-2-oxovaleric acid. These decreased in *B. thetaiotaomicron* Δcps after DAC15 exposure, with a reduction of 3.1- and 5.1-fold change respectively (**Fig. 4A, B**). These metabolites are catabolic products of isoleucine and leucine produced during branched-chain amino acid degradation, which suggests that modulation of this metabolic pathway may occur in response to phage exposure in this strain. This alteration in the metabolism of iso- and leucine catabolic products could indicate a metabolic shift aimed at redirecting resources or energy metabolism in response to the stress induced by DAC15 infection itself, or a population-level biosynthetic response to it (production of additional defensive cell surface structures). No significant differences were detected among the metabolites of the other tested strains.

**Fig. 4.**
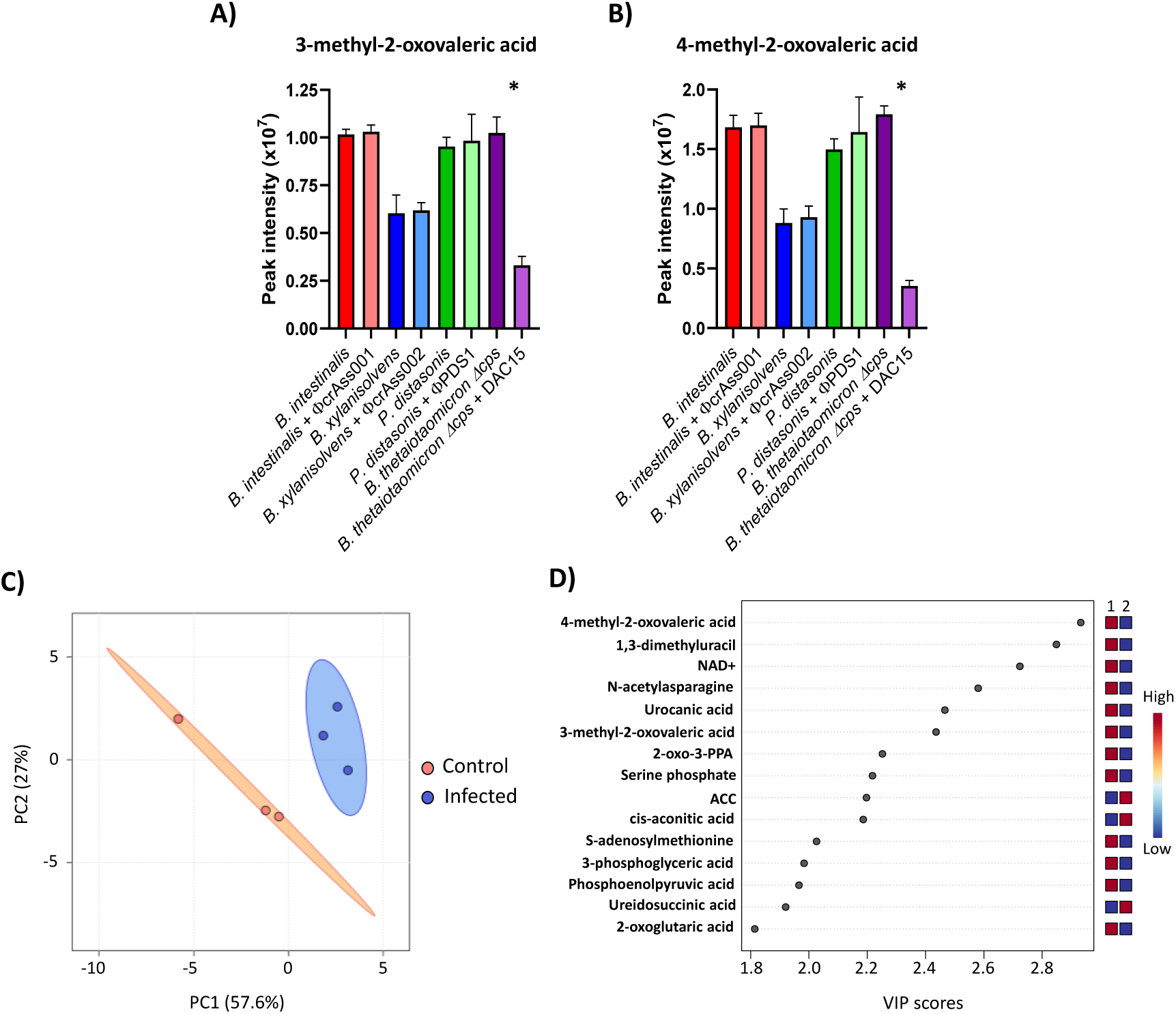
Intracellular metabolomic changes in *B. thetaiotaomicron* Δcps after DAC15 infection. **(A-B)** LC-MS peak intensities of 3-methyl-2-oxovaleric acid **(A)** and 4-methyl-2-oxovaleric acid **(B)** for each phage-host pair, obtained from untargeted intracellular metabolomics analysis performed in the samples obtained during mid-log phase growth on the fifth day of the co-culture experiment. Bars represent the mean of triplicate biological samples, with error bars indicating the standard deviation (SD). * Significant differences were set up at *p* < 0.05. **(C)** Principal Component Analysis (PCA) of intracellular metabolomic profiles from *B. thetaiotaomicron* Δcps and DAC15 co-cultures, based on detected compounds with accuracy levels 1 and 2a, extracted from bacterial pellets. Orange indicates the control condition (uninfected), while blue is the infected condition. Ellipses represent 95% confidence intervals, and the explained variances of the two principal components are shown in brackets along the axes. **(D)** Important features identified by partial least squares-discriminant analysis (PLS-DA) in *B. thetaiotaomicron* Δcps bacterial pellets collected at the mid-log phase on the fifth day of the co-culture. The coloured boxes on the right indicate the relative peak area of each corresponding metabolite in the groups under study, with red indicating higher peak areas and blue indicating lower. Group 1 corresponds to the control condition, and group 2 corresponds to the DAC-15 infected samples. 2-oxo-3-PPA: 2-oxo-3-phenylpropanoic acid; ACC: 1-Aminocyclopropanecarboxylic acid.

A multivariate analysis was performed to investigate global differences in the metabolome of these strains after phage exposure. PCA revealed the separation of individual samples of *B. thetaiotaomicron* Δcps driven by phage infection (**Fig. 4C**). The total variance explained by the two principal components (PCs) was 84.6% (PC1 = 57.6% and PC2 = 27%). This clear grouping was not observed for the samples of the capsular strains (**Fig. S8**). Although a tendency to be grouped could be observed in *B. intestinalis* and *P. distasonis* metabolome samples depending on whether they were infected or not, the small number of replicates did not provide sufficient statistical power.

Next, a partial least square discriminant analysis (PLS-DA), a supervised method with full awareness of the class labels, was applied. The metabolome samples were grouped according to whether the cultures were infected or not in all cases. However, only the model for *B. thetaiotaomicron* Δcps showed good model fit, quality, and accuracy after cross-validation (R2 = 0.97, Q2 = 0.82), which confirmed that significant changes occurred in the metabolome of this strain after infection with DAC15 (**Table S8**). Variable Importance in Prediction (VIP) scores, which estimate the contribution of a given predictor to a PLS regression model, were calculated and are shown in **Fig. 4D** and **Table S9**. The compounds that contributed most strongly to the separation between the sample groups in *B. thetaiotaomicron* Δcps were 4-methyl-2-oxovaleric acid, 1,3-dimethyluracil, NAD+, N-acetylasparagine, urocanic acid, 3-methyl-2-oxovaleric acid, 2-oxo-3-phenylpropanoic acid, serine phosphate, S-adenosylmethionine (decreased in infection), and 1-aminocyclopropanecarboxylic acid and cis-aconitic acid (increased in infection). Most of these metabolites are related to amino acid pathways, which suggests that the *B. thetaiotaomicron* Δcps population responds to DAC15 infection with metabolic changes by trying to modify different metabolic pathways to achieve survival despite the phage attack.

In the case of capsular strains, the cross-validation of these models showed poor model fit and accuracy (**Table S8**). These findings, along with our previous results, confirm that the metabolites detected using this approach may not be sufficiently relevant to explain significant changes in the metabolome of these strains. Focusing this metabolomics analysis on polar metabolites could yield more relevant results for these strains, especially considering the critical role of sugars in producing CPSs.

## 4. Discussion

Despite being constantly exposed to external perturbing factors, such as dietary changes, antibiotic therapy or infectious diseases, the gut microbiome is characterised by its resilience to perturbations and its high temporal stability [48,49]. This stability has also been reported for the gut virome together with a pronounced interindividual variability, with virulent crAss-like phages being one of the most prevalent entities in the human gut virome capable of long-term co-existence with their Bacteroidales hosts [23,50]. Here, we focused on four lytic gut phages infecting bacteria of the order Bacteroidales to identify possible new mechanisms for this co-existence. Two distinct phylogenetic phage groups were included in our analysis: crAss-like phages ϕcrAss001 - *B. intestinalis*, ϕcrAss002 - *B.xylanisolvens*, DAC15 - *B.thetaiotaomicron*, and siphovirus-like phage ϕPDS1 - *P. distasonis*, as well as hosts with two different phenotypes (capsular and acapsular).

The expectation is that in a simple monoculture model like ours, lytic phages should exert a strong top-down control on the host population density favouring the emergence of resistant mutants. These mutants can completely or partially replace the ancestral sensitive population, depending on the cost of resistance, which in turn leads to the reduction or complete extinction of the phage population [51]. Our co-culture experiment revealed that phage persistence coincides with the co-existence of sensitive and resistant bacterial populations in an equilibrium (mainly due to phase variation of surface antigens), allowing bacterial survival due to the resistant populations, and phage propagation due to the sensitive populations. The initially low percentage of phage-resistant host cells increased to over 90% when in contact with the phages. A 100% resistant population would not be expected given that the phages continued to propagate, so some of our results showing values close to 100% (**Fig.1E, F, G, H**) are explained by the limitations of plating assays.

The observation of stable co-existence in our phage-bacteria pairs can be linked with the eco-evolutionary model of fluctuating selection dynamics, where density-dependent fluctuating selection operates through a trade-off between the benefits of resistance and its associated metabolic costs [10,13], promoting prolonged co-existence of phage and host. Phage ecology is highly variable across different environmental microbiomes. For instance, in the ocean, phage-host pair dynamics are best explained by the ‘kill-the-winner’ model, with 20-40% of ocean bacteria being lysed by phage every day [52]. In contrast, in soil microbiomes, lysogeny is favoured, with ecological dynamics being best explained by the ‘piggyback the winner’ model. Under this model, phages lysogenize their hosts during periods of high microbial abundance and growth rates, allowing them to multiply in parallel with their host [52,53]. An extension of this model, accounting for the absence of true lysogeny, could also explain some of the behaviour observed in our study. Recently, the existence of phage P1-like vegetative replication without producing phage progeny has been proposed for the prototypical crAssphage – the founding species in the order *Crassvirales* [54]. It is possible that phages like prototypical crAssphage and crAss002, both incapable of forming plaques in their host cultures, may instead form transient pseudo-lysogens by replicating their genomes via this vegetative replication pathway. Thus, a combination of fluctuating selection dynamics (dynamic equilibrium between sensitive and resistant subpopulations) with ‘piggyback-the-winner’ dynamics (transient pseudolysogeny) could in theory be occurring in these models. Different eco-evolutionary models might be useful in describing certain aspects of phage-host co-existence in complex microbial communities such as the human gut microbiome [52]. However, even the very simplistic single pair models used in this study highlight the complexity of phage-host interactions and the inadequacy of a simple model to capture them [13]. The relevance of correctly capturing these dynamics lies in their potential involvement in microbiome transformation in human gut diseases such as inflammatory bowel disease, where bacterial phase variations driven by phages and host inflammation have been described to contribute to bacterial functional plasticity during disease [55].

One characteristic of gut Bacteroidales is their ability to alter their surfaces dynamically to gain a survival advantage in their ecosystem, protecting themselves from host and environmental factors [56]. This includes phase variation (expression of alternative types) of CPS driven by invertible promoters alternating between “ON” and “OFF” orientations [57,58]. On top of that, NusG-like anti-termination factors [59], and trans locus inhibitors [42] were proposed to ensure the expression of only one type of CPS at a time in any given cell. This mechanism has been reported to modify phage susceptibility and allow for prolonged phage persistence in some strains of *B. intestinalis* and *B. thetaiotaomicron* [22,24]. Our transcriptomics analysis confirms this previously proposed model and extends it to two new phage-host pairs in *B. xylanisolvens* and *P. distasonis*, where operons associated with alternative CPS types showed the strongest transcriptional responses, with some being upregulated (protective) and others downregulated (permissive) (**Fig. 2A, B, C, D**). Specifically, our results for *B. intestinalis* are consistent with those obtained in Shkoporov et al. (2019) [22], which reported strong repression of the PVR9 CPS operon and the upregulation of CPS operons associated with loci PVR7, PVR8 and PVR11 during the stationary phase of a phage-bacteria co-culture. All these loci share an IR sequence (GTTCGTTTAA), flanking their putative invertible promoters. The only difference from this previous study is that PVR12, which also shares this promoter sequence, has been upregulated in our study, showing a stronger response than PVR8 and PVR11, while Shkoporov et al. (2019) [22] did not observe any significant change for this locus. This difference could be attributed to the different timing of sample collection in the experiments, as well as the stochasticity of phase variation and the following selection of phage-resistant variants. When infected by their corresponding phages, *B. xylanisolvens* and *P. distasonis* showed down- and up-regulation of CPS operons, mediated by invertible promoters flanked by conserved IR sequences (GTTACTTCTTAGGTAACGGA for *B. xylanisolvens* and GCTACTYRGNRAGTAGC for *P. distasonis*). One remarkable difference between these IR sequences with respect to the one from *B. intestinalis* is that they contain internal palindromic regions and are preceded by integrase or invertase enzymes, which suggests that while the overall mechanism is broadly similar, specific variations are present. When compared to other *Bacteroides* species, such as *B. thetaiotaomicron* or *B. fragilis* [24,45], these IR sequences are more similar and even share a common pattern involving a short fragment (GTTAC), with *P. distasonis* replacing the first thymine with a cytosine. This indicates that the *B. intestinalis* system differs more from those of the other strains. Additionally, these palindromic IR sequences were found not only near CPS operons but also in other genomic regions, where they were also preceded by recombinases, indicating a conserved regulatory mechanism dependent on site-specific recombination.

Common elements are found at the beginning of most of these CPS loci, including genes encoding proteins of the UpxY and UpxZ families [42,59]. While UpxY-like proteins positively regulate transcription of their respective CPS biosynthesis operon by preventing premature transcription termination in the untranslated region [59], UpxZ-like proteins repress transcription of non-cognate CPS [42]. In the case of *P. distasonis*, only two of the twelve CPS loci have this UpxY family of proteins annotated in its genome. The rest of the CPS loci are annotated as transcriptional regulators – anti-termination protein NusG. Although these transcriptional factors may correspond to the UpxY system, as genomic analysis of other *P. distasonis* strains has shown [60], other NusG-mediated mechanisms could also be attributed. This family of proteins is usually preceded by recombinases and invertible promoters, which makes these regions hotspots for recombination and enables an ON/OFF switching mechanism that continuously shifts, influencing the permissiveness to phage infection. Gene clusters downregulated during phage infection may increase susceptibility to phages, while those that were upregulated could have a protective or neutral function. This dynamic equilibrium between resistant and sensitive bacterial sub-populations helps maintain phage persistence. However, this is not the only mechanism utilised by Bacteroidales to facilitate phage persistence, as evidenced by our observations during the propagation of the crAss-phage DAC15 in an acapsular strain. A diverse transcriptional response after DAC15 infection was observed in the knockout strain. BT_RS09760 (also named BT1927) was the most highly upregulated gene, which is located near a site-specific integrase and has been reported to encode an S-layer protein homologue, one of several related by distinct homologues encoded in *B. thetaiotaomicron* genome, all of which are expressed in a phase-variable manner [46]. The authors described that in a wildtype *B. thetaiotaomicron* culture, the expression of BT1927 was very low (∼1:1,000 cells), and when expressed, it increased bacterial resistance to complement-mediated killing. Our results reflect a strong upregulation after DAC15 infection, with proteomics supporting significant differences in this protein in both the cell pellet and the culture supernatant. This suggests an alternative adaptative mechanism of *Bacteroides* to phage exposure when CPSs are absent, which is also related to phase variation. Another study also observed this upregulation when the strain was infected with a different phage, ARB25 [24]. Similarly to the mechanism of CPS phase variation, the S-layer operons are also preceded by promoters flanked by consensus IR sequences (CCGTTACCTABVRAGTAAC), which were detected in other 7 upregulated loci next to site-specific integrases S-layer surface proteins and OmpA family proteins, as previously reported [24]. This IR sequence also has a small internal repeat with the common element CGTTA shared with the IR sequences of the CPS operons in the other *Bacteroides* species previously discussed. In fact, four of the CPS operons (CPS1-BT_RS01830, CPS3-BT_RS02920, CPS5-BT_RS08400, CP6-BT_RS08750) in *B. thetaiotaomicron* wildtype contain a nearly identical IR sequence (CCGTTACCTAAGAAGTAAC) flanking their invertible promoter and neighboured by tyrosine-type DNA invertases. This finding shows that the mechanism of phase variation of these different cell surface operons might be similarly regulated, supporting phage persistence even in the absence of CPS. The highly diverse transcriptional responses observed when alternative CPS cannot be switched imply the existence of multiple layers of secondary phage resistance mechanisms, each of which being perhaps less efficient than CPS switching, but together, they can produce a sufficient quantitative effect to allow the population to survive in the presence of phage. This can be justified by the longer time required for this strain to achieve a high percentage of resistant cells compared to the capsulated strains (**Fig. 1E, F, G, H**).

The differential expression of alternative CPS types may also cause some intermediate metabolism changes in the bacterial population under attack from phage. Here, we analysed the potential metabolic changes associated with persistent phage infection in populations of *Bacteroides spp*. host strains. Previous studies that have investigated changes in the metabolome profile caused by phage predation have been conducted with *in vivo* models, focusing their attention on how it affects the gut microbiome composition and the gut metabolome of mammalian hosts [61,62], rather than on the impact on the bacterial host. The acapsular strain of *B. thetaiotaomicron* showed the highest differences in its intracellular metabolome. Most of these metabolites were related to amino acid metabolism, involving isoleucine, leucine, phenylalanine and histidine amino acids. This was reflected in the reduction of abundances of 3-methyl-2-oxovaleric acid, 4-methyl-2-oxovaleric acid, 2-oxo-3-phenylpropanoic acid, and urocanic acid, which are intermediate compounds in the degradation pathways of these amino acids [63, 64]. This reduction may indicate that the metabolic pathways for amino acid degradation, which can be used as a source of energy for bacterial growth, can be reduced due to the presence of phage. Another relevant change was the depletion of NAD+ when DAC15 infected the host, an essential metabolite for numerous metabolic processes, acting as a cofactor in cellular metabolism and redox reactions. This reduction of NAD+ has been described as a defence mechanism used against phage infection [7,65,66,67], thus depriving the phage of this molecule and impeding phage propagation. It appears that the absence of CPS forces bacteria to use other mechanisms to defend against phage, which may explain these changes. However, the capsulated strains showed fewer changes in their metabolome, possibly mediated by the ability to deploy a less costly type of mechanism for phage resistance – the alternative CPS types. Also, the technique used for this analysis was focused on semi-polar compounds. This may have resulted in certain compounds not being extracted, such as those with high polarity, including polar sugars serving as intermediates in CPS biosynthesis.

## 5. Conclusions

This study represents an additional step in our understanding of the mechanisms of the long-term persistence of virulent bacteriophages in their host cultures, including the persistence of crAss-like and non-crAss phages in *Bacteroides* and *Parabacteroides*. Phase-variable CPS expression plays an essential role in the bacterial ability to co-exist with phages. Other gene products, including but not limited to the alternative types of S-layer proteins, regulated by phase variation also appear to enable efficient co-existence with phages, suggesting cooperative relationships between different mechanisms of resistance. In the absence of CPS, higher transcriptomic, proteomic, and metabolomic changes are observed since more factors are involved in achieving the equilibrium between bacterial and host populations. Further research is necessary to delve deeper into these aspects, enhancing our understanding of phage-bacteria interactions in the crowded environment of the gut.

## Availability of data and material

All data generated or analysed during this study are included in this published paper and its supplementary material. Raw sequencing data for transcriptomics are available from NCBI Sequence Read Archive (SRA) repository under BioProject PRJNA1131275. Proteomics data including raw mass spectrometry files and databases are available from ProteomeXchange Consortium via the PRIDE [68] partner repository with the dataset identifier PXD057336 [Reviewer Access at: https://www.ebi.ac.uk/pride/login with Reviewer Username: Password: ICAzyG65Y9kx].

## Author contributions

Conceptualization: A.N.S. and C.H.; Methodology: A.C.M, A.N.S, C.H., M.K. and R.P.R.; Investigation: A.C.M, C.B., J.L.M. and C.A.T.; Formal Analysis: A.C.M, C.B. and J.L.M.; Writing— original draft: A.C.M.; Writing—review and editing: A.C.M., A.N.S., C.H. and C.B.; Supervision: A.N.S., C.H. and M.K.; Project administration: L.A.D.; Funding acquisition: A.C.M., A.N.S., C.H., M.K. and R.P.R. All authors read and approved the final manuscript.

## Acknowledgements

We thank Rachel Moser for technical support with proteomics. We made all LC-MS/MS measurements in the Molecular Education, Technology, and Research Innovation Center (METRIC) at North Carolina State University.

## References

1. Fan Y, Pedersen O. Gut microbiota in human metabolic health and disease. Nat Rev Microbiol. 2021;19(1):55–71.

2. de Vos WM, Tilg H, Van Hul M, Cani PD. Gut microbiome and health: mechanistic insights. Gut. 2022;71(5):1020–1032.

3. Shareefdeen H, Hill C. The gut virome in health and disease: new insights and associations. Curr Opin Gastroenterol. 2022;38(6):549–554.

4. Tobin CA, Hill C, Shkoporov AN. Factors Affecting Variation of the Human Gut Phageome. Annu Rev Microbiol. 2023;77:363–379.

5. Shkoporov AN, Turkington CJ, Hill C. Mutualistic interplay between bacteriophages and bacteria in the human gut. Nat Rev Microbiol. 2022;20(12):737–749.

6. Piel D, Bruto M, Labreuche Y, Blanquart F, Goudenège D, Barcia-Cruz R, et al. Phage-host coevolution in natural populations. Nat Microbiol. 2022;7(7):1075–1086.

7. Millman A, Melamed S, Leavitt A, Doron S, Bernheim A, Hör J, et al. An expanded arsenal of immune systems that protect bacteria from phages. Cell Host Microbe. 2022;30(11):1556–1569.e5.

8. Chevallereau A, Pons BJ, van Houte S, Westra ER. Interactions between bacterial and phage communities in natural environments. Nat Rev Microbiol. 2022;20(1):49–62.

9. Koskella B, Brockhurst MA. Bacteria-phage coevolution as a driver of ecological and evolutionary processes in microbial communities. FEMS Microbiol Rev. 2014;38(5):916–31.

10. Scanlan PD. Bacteria-Bacteriophage Coevolution in the Human Gut: Implications for Microbial Diversity and Functionality. Trends Microbiol. 2017;25(8):614–623.

11. De Sordi L, Lourenço M, Debarbieux L. “I will survive“: A tale of bacteriophage-bacteria coevolution in the gut. Gut Microbes. 2019;10(1):92–99.

12. Shkoporov AN, Hill C. Bacteriophages of the Human Gut: The “Known Unknown” of the Microbiome. Cell Host Microbe. 2019;25(2):195–209.

13. Castledine M, Buckling A. Critically evaluating the relative importance of phage in shaping microbial community composition. Trends Microbiol. 2024;32(10):957–969.

14. Mirzaei MK, Maurice CF. *Ménage à trois* in the human gut: interactions between host, bacteria and phages. Nat Rev Microbiol. 2017;15(7):397–408.

15. Smith L, Goldobina E, Govi B, Shkoporov AN. Bacteriophages of the Order *Crassvirales*: What Do We Currently Know about This Keystone Component of the Human Gut Virome? Biomolecules. 2023;13(4):584.

16. Wexler AG, Goodman AL. An insider’s perspective: *Bacteroides* as a window into the microbiome. Nat Microbiol. 2017;2:17026.

17. Shkoporov AN, Khokhlova EV, Fitzgerald CB, Stockdale SR, Draper LA, Ross RP, et al. ΦCrAss001 represents the most abundant bacteriophage family in the human gut and infects *Bacteroides intestinalis*. Nat Commun. 2018;9(1):4781.

18. Hryckowian AJ, Merrill BD, Porter NT, Van Treuren W, Nelson EJ, Garlena RA, et al. *Bacteroides thetaiotaomicron*-Infecting Bacteriophage Isolates Inform Sequence-Based Host Range Predictions. Cell Host Microbe. 2020;28(3):371–379.e5.

19. Guerin E, Shkoporov AN, Stockdale SR, Comas JC, Khokhlova EV, Clooney AG, et al. Isolation and characterisation of ΦcrAss002, a crAss-like phage from the human gut that infects *Bacteroides xylanisolvens*. Microbiome. 2021;9(1):89.

20. Ramos-Barbero MD, Gómez-Gómez C, Sala-Comorera L, Rodríguez-Rubio L, Morales-Cortes S, Mendoza-Barberá E, et al. Characterization of crAss-like phage isolates highlights *Crassvirales* genetic heterogeneity and worldwide distribution. Nat Commun. 2023;14(1):4295.

21. Papudeshi B, Vega AA, Souza C, Giles SK, Mallawaarachchi V, Roach MJ, et al. Host interactions of novel *Crassvirales* species belonging to multiple families infecting bacterial host, *Bacteroides cellulosilyticus* WH2. Microb Genom. 2023;9(9):001100.

22. Shkoporov AN, Khokhlova EV, Stephens N, Hueston C, Seymour S, Hryckowian AJ, et al. Long-term persistence of crAss-like phage crAss001 is associated with phase variation in *Bacteroides intestinalis*. BMC Biol. 2021;19(1):163.

23. Shkoporov AN, Clooney AG, Sutton TDS, Ryan FJ, Daly KM, Nolan JA, et al. The Human Gut Virome Is Highly Diverse, Stable, and Individual Specific. Cell Host Microbe. 2019;26(4):527–541.e5.

24. Porter NT, Hryckowian AJ, Merrill BD, Fuentes JJ, Gardner JO, Glowacki RWP, et al. Phase-variable capsular polysaccharides and lipoproteins modify bacteriophage susceptibility in *Bacteroides thetaiotaomicron*. Nat Microbiol. 2020;5(9):1170–1181.

25. Li J, Feng S, Yu L, Zhao J, Tian F, Chen W, et al. Capsular polysaccarides of probiotics and their immunomodulatory roles. Food Sci. Hum. Wellness. 2022;11(5):1111–1120.

26. Cortés-Martín A, Denise R, Guerin E, Stockdale SR, Draper LA, Ross RP, et al. Isolation and characterization of a novel lytic *Parabacteroides distasonis* bacteriophage φPDS1 from the human gut. Gut Microbes. 2024;16(1):2298254.

27. Love MI, Anders S, Kim V, Huber W. RNA-Seq workflow: gene-level exploratory analysis and differential expression. F1000Res. 2015;4:1070.

28. Langmead B, Salzberg SL. Fast gapped-read alignment with Bowtie 2. Nat Methods. 2012;9(4):357–359.

29. Li H. New strategies to improve minimap2 alignment accuracy. Bioinformatics. 2021;37(23):4572–4574.

30. Sedlazeck FJ, Rescheneder P, Smolka M, Fang H, Nattestad M, von Haeseler A, et al. Accurate detection of complex structural variations using single-molecule sequencing. Nat Methods. 2018;15(6):461–468.

31. Danecek P, Bonfield JK, Liddle J, Marshall J, Ohan V, Pollard MO, et al. Twelve years of SAMtools and BCFtools. Gigascience. 2021;10(2):giab008.

32. Bolyen E, Rideout JR, Dillon MR, Bokulich NA, Abnet CC, Al-Ghalith GA, et al. Reproducible, interactive, scalable and extensible microbiome data science using QIIME 2. Nat Biotechnol. 2019;37(8):852–857.

33. Tatusov RL, Galperin MY, Natale DA, Koonin EV. The COG database: a tool for genome-scale analysis of protein functions and evolution. Nucleic Acids Res. 2000;28(1):33–36.

34. Camacho C, Coulouris G, Avagyan V, Ma N, Papadopoulos J, Bealer K, et al. BLAST+: architecture and applications. BMC Bioinformatics. 2009;10:421.

35. Gilchrist CLM, Chooi YH. clinker & clustermap.js: automatic generation of gene cluster comparison figures. Bioinformatics. 2021;37(16):2473–2475.

36. Beyer NH, Schou C, Houen G, Heegaard NH. Extraction and identification of electroimmunoprecipitated proteins from agarose gels. J Immunol Methods. 2008; 330(1-2):24–33.

37. Oberg AL, Vitek O. Statistical design of quantitative mass spectrometry-based proteomic experiments. J Proteome Res. 2009;8(5):2144–56.

38. Mordant A, Kleiner M. Evaluation of Sample Preservation and Storage Methods for Metaproteomics Analysis of Intestinal Microbiomes. Microbiol Spectr. 2021;9(3):e0187721.

39. Blakeley-Ruiz JA, Bartlett A, McMillan AS, Awan A, Vanhoy Walsh M, Meyerhoffer AK, et al. Dietary protein source strongly alters gut microbiota composition and function. bioRxiv [Preprint]. 2024;2024.04.04.588169.

40. Doneanu CE, Chen W, Mazzeo JR. UPLC/MS Monitoring of Water-Soluble Vitamin Bs in Cell Culture Media in Mins. 2011. Waters. https://www.waters.com/webassets/cms/library/docs/720004042en.pdf (2011).

41. Pang Z, Lu Y, Zhou G, Hui F, Xu L, Viau C, et al. MetaboAnalyst 6.0: towards a unified platform for metabolomics data processing, analysis and interpretation. Nucleic Acids Res. 2024;52(W1):W398–W406.

42. Chatzidaki-Livanis M, Weinacht KG, Comstock LE. Trans locus inhibitors limit concomitant polysaccharide synthesis in the human gut symbiont *Bacteroides fragilis*. Proc Natl Acad Sci U S A. 2010;107(26):11976–11980.

43. Saba J, Flores K, Marshall B, Engstrom MD, Peng Y, Garje AS, et al. *Bacteroides* expand the functional versatility of a universal transcription factor and transcribed DNA to program capsule diversity. bioRxiv [Preprint]. 2024;2024.06.21.599965.

44. Grondin JM, Tamura K, Déjean G, Abbott DW, Brumer H. Polysaccharide Utilization Loci: Fueling Microbial Communities. J Bacteriol. 2017;199(15):e00860–16.

45. Weinacht KG, Roche H, Krinos CM, Coyne MJ, Parkhill J, Comstock LE. Tyrosine site-specific recombinases mediate DNA inversions affecting the expression of outer surface proteins of *Bacteroides fragilis*. Mol Microbiol. 2004;53(5):1319–1330.

46. Taketani M, Donia MS, Jacobson AN, Lambris JD, Fischbach MA. A Phase-Variable Surface Layer from the Gut Symbiont *Bacteroides thetaiotaomicron*. mBio. 2015;6(5):e01339–15.

47. Lan F, Saba J, Qian Y, Ross T, Landick R, Venturelli OS. Single-cell analysis of multiple invertible promoters reveals differential inversion rates as a strong determinant of bacterial population heterogeneity. Sci Adv. 2023;9(31):eadg5476.

48. Mehta RS, Abu-Ali GS, Drew DA, Lloyd-Price J, Subramanian A, Lochhead P, et al. Stability of the human faecal microbiome in a cohort of adult men. Nat Microbiol. 2018;3(3):347–355.

49. Fassarella M, Blaak EE, Penders J, Nauta A, Smidt H, Zoetendal EG. Gut microbiome stability and resilience: elucidating the response to perturbations in order to modulate gut health. Gut. 2021;70(3):595–605.

50. Garmaeva S, Gulyaeva A, Sinha T, Shkoporov AN, Clooney AG, Stockdale SR, et al. Stability of the human gut virome and effect of gluten-free diet. Cell Rep. 2021;35(7):109132.

51. Bohannan BJM, Lenski RE. Linking Genetic Change to Community Evolution: Insights from Studies of Bacteria and Bacteriophage. Ecol. Lett. 2000;3:362–377.

52. Brown TL, Charity OJ, Adriaenssens EM. Ecological and functional roles of bacteriophages in contrasting environments: marine, terrestrial and human gut. Curr Opin Microbiol. 2022;70:102229.

53. Silveira CB, Rohwer FL. Piggyback-the-Winner in host-associated microbial communities. NPJ Biofilms Microbiomes. 2016;2:16010.

54. Schmidtke DT, Hickey AS, Liachko I, Sherlock G, Bhatt AS. Analysis and culturing of the prototypic crAssphage reveals a phage-plasmid lifestyle. bioRxiv [Preprint]. 2024;2024.03.20.585998.

55. Carasso S, Zaatry R, Hajjo H, Kadosh-Kariti D, Ben-Assa N, Naddaf R, et al. Inflammation and bacteriophages affect DNA inversion states and functionality of the gut microbiota. Cell Host Microbe. 2024;32(3):322–334.e9.

56. Coyne MJ, Comstock LE. Niche-specific features of the intestinal bacteroidales. J Bacteriol. 2008;190(2):736–742.

57. Coyne MJ, Weinacht KG, Krinos CM, Comstock LE. Mpi recombinase globally modulates the surface architecture of a human commensal bacterium. Proc Natl Acad Sci U S A. 2003;100(18):10446–10451.

58. Hickey CA, Kuhn KA, Donermeyer DL, Porter NT, Jin C, Cameron EA, et al. Colitogenic *Bacteroides thetaiotaomicron* Antigens Access Host Immune Cells in a Sulfatase-Dependent Manner via Outer Membrane Vesicles. Cell Host Microbe. 2015;17(5):672–680.

59. Chatzidaki-Livanis M, Coyne MJ, Comstock LE. A family of transcriptional antitermination factors necessary for synthesis of the capsular polysaccharides of *Bacteroides fragilis*. J Bacteriol. 2009;191(23):7288–7295.

60. Chamarande J, Cunat L, Alauzet C, Cailliez-Grimal C. *In Silico* Study of Cell Surface Structures of *Parabacteroides distasonis* Involved in Its Maintenance within the Gut Microbiota. Int J Mol Sci. 2022;23(16):9411.

61. Hsu BB, Gibson TE, Yeliseyev V, Liu Q, Lyon L, Bry L, et al. Dynamic Modulation of the Gut Microbiota and Metabolome by Bacteriophages in a Mouse Model. Cell Host Microbe. 2019;25(6):803–814.e5.

62. Lorenzo-Rebenaque L, Casto-Rebollo C, Diretto G, Frusciante S, Rodríguez JC, Ventero MP, et al. Examining the effects of *Salmonella* phage on the caecal microbiota and metabolome features in *Salmonella*-free broilers. Front Genet. 2022;13:1060713.

63. Tang CM, Lin G, Chiang MH, Yeh KW, Huang JL, Su KW, et al. Longitudinal Metabolomic Analysis Reveals Gut Microbial-Derived Metabolites Related to Formula Feeding and Milk Sensitization Development in Infancy. Metabolites. 2022;12(2):127.

64. Zhu Y, Dwidar M, Nemet I, Buffa JA, Sangwan N, Li XS, et al. Two distinct gut microbial pathways contribute to meta-organismal production of phenylacetylglutamine with links to cardiovascular disease. Cell Host Microbe. 2023;31(1):18–32.e9.

65. Garb J, Lopatina A, Bernheim A, Zaremba M, Siksnys V, Melamed S, et al. Multiple phage resistance systems inhibit infection via SIR2-dependent NAD+ depletion. Nat Microbiol. 2022;7(11):1849–1856.

66. Ofir G, Herbst E, Baroz M, Cohen D, Millman A, Doron S, et al. Antiviral activity of bacterial TIR domains via immune signalling molecules. Nature. 2021;600(7887):116–120.

67. Yin H, Li X, Wang X, Zhang C, Gao J, Yu G, et al. Insights into the modulation of bacterial NADase activity by phage proteins. Nat Commun. 2024;15(1):2692.

68. Perez-Riverol Y, Bai J, Bandla C, García-Seisdedos D, Hewapathirana S, Kamatchinathan S, et al. The PRIDE database resources in 2022: a hub for mass spectrometry-based proteomics evidences. Nucleic Acids Res. 2022;50(D1):D543–D552.

