## Supplementary Figures for "Adaptations in gut Bacteroidales facilitate stable co-existence with their lytic bacteriophages"

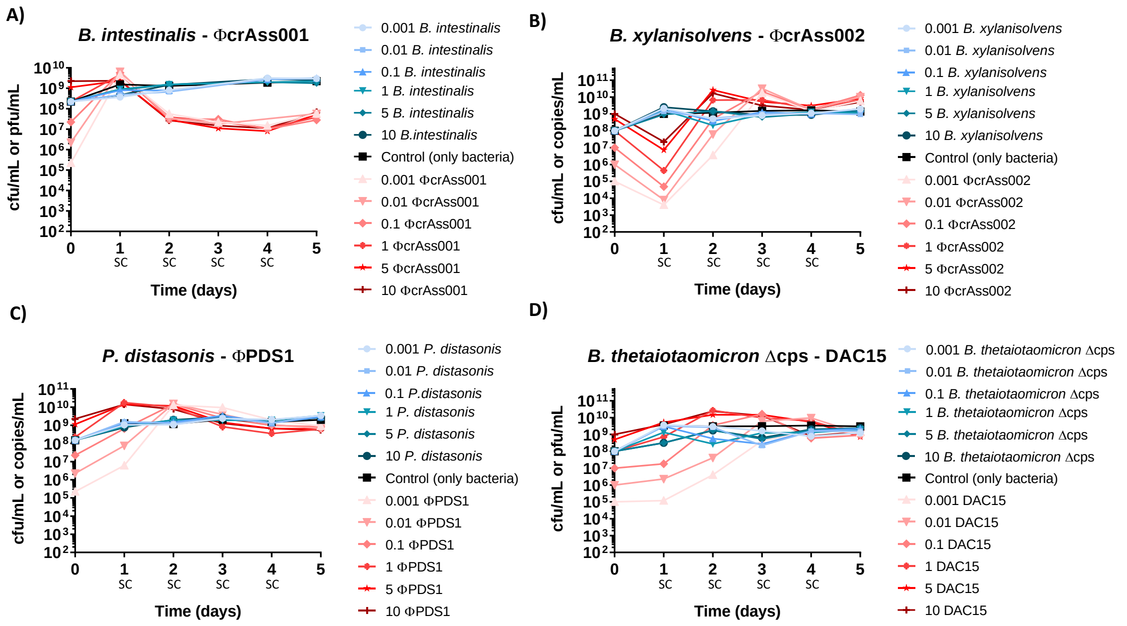


**Fig. S1.** Enumeration of bacteria and phage during the 5-day co-culture experiment testing a range of multiplicities of infection (MOIs) of phage (0.001 – 10) at the start of the test. After each 24-hour period, a new subculture (SC) was performed in fresh media at a dilution of 1:50. **(A)** *B. intestinalis* + ϕcrAss001, **(B)** *B. xylanisolvens* + ϕcrAss002, **(C)** *P. distasonis* + ϕPDS1, **(D)** *B. thetaiotaomicron* Δcps + DAC15. The black line represents bacterial counts (CFU/mL) for the control condition (uninfected bacteria), the blue lines represent bacterial counts (CFU/mL) in co-culture with phage at different MOIs (lighter lines indicate lower MOIs), and the red lines represent the phage counts (pfu/mL or copies/mL) in co-culture with bacteria at different MOIs (lighter lines indicate lower MOIs). Differences in phage enumeration methods were due to the ability to produce countable plaques (CFU/mL) or, alternatively, qPCR (copies/mL) when plaques were not detected.

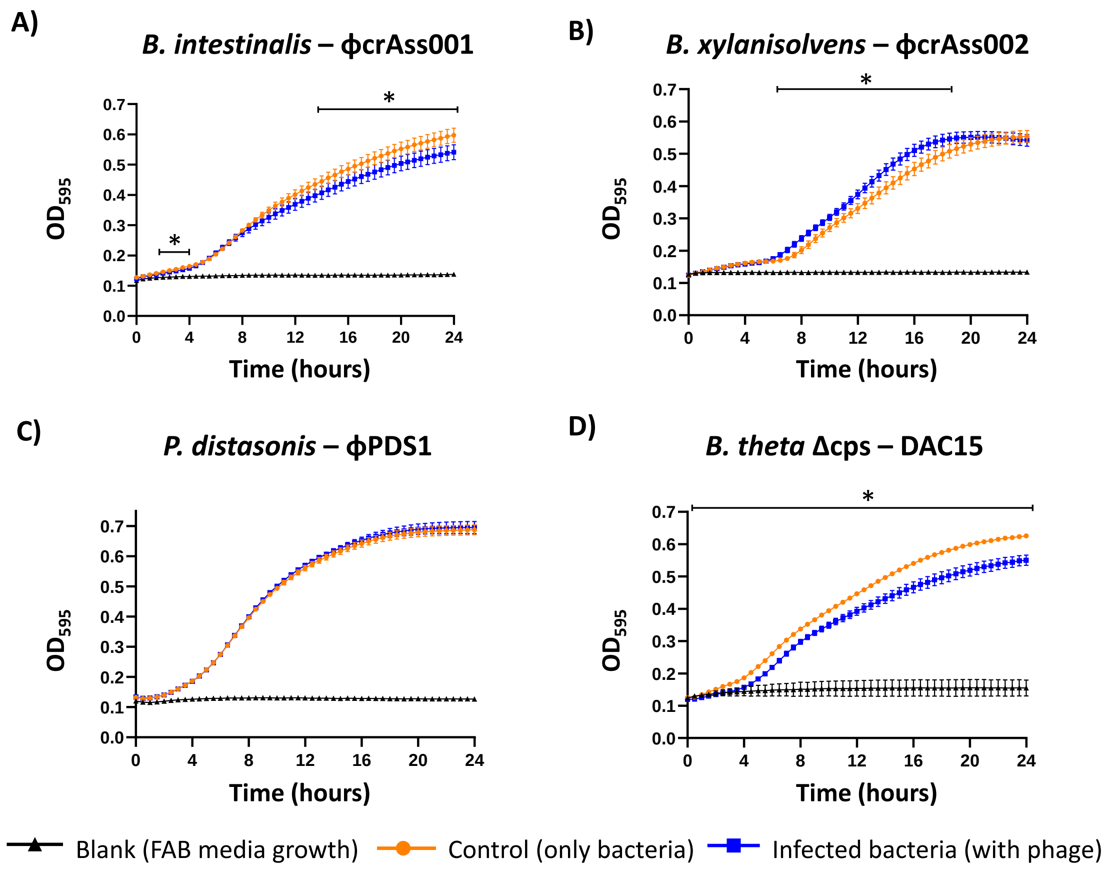


**Fig. S2.** Kinetics of bacterial growth during 24 hours after 5 days of co-culture, measured by optical density ($OD_{595}$). **(A)** *B. intestinalis*, **(B)** *B. xylanisolvens*, **(C)** *P. distasonis*, **(D)** *B. thetaiotaomicron* Δcps. The orange line represents $OD_{595}$ of control bacterial cultures after 5 days of subculturing, the blue line represents $OD_{595}$ of infected bacterial cultures after 5 days of subculturing, and the black line represents the blank (growth media only, FAB). Measurements were carried out in triplicate. *: Significant differences in $OD_{595}$ when comparing the control with the infected condition determined at $p < 0.05$.

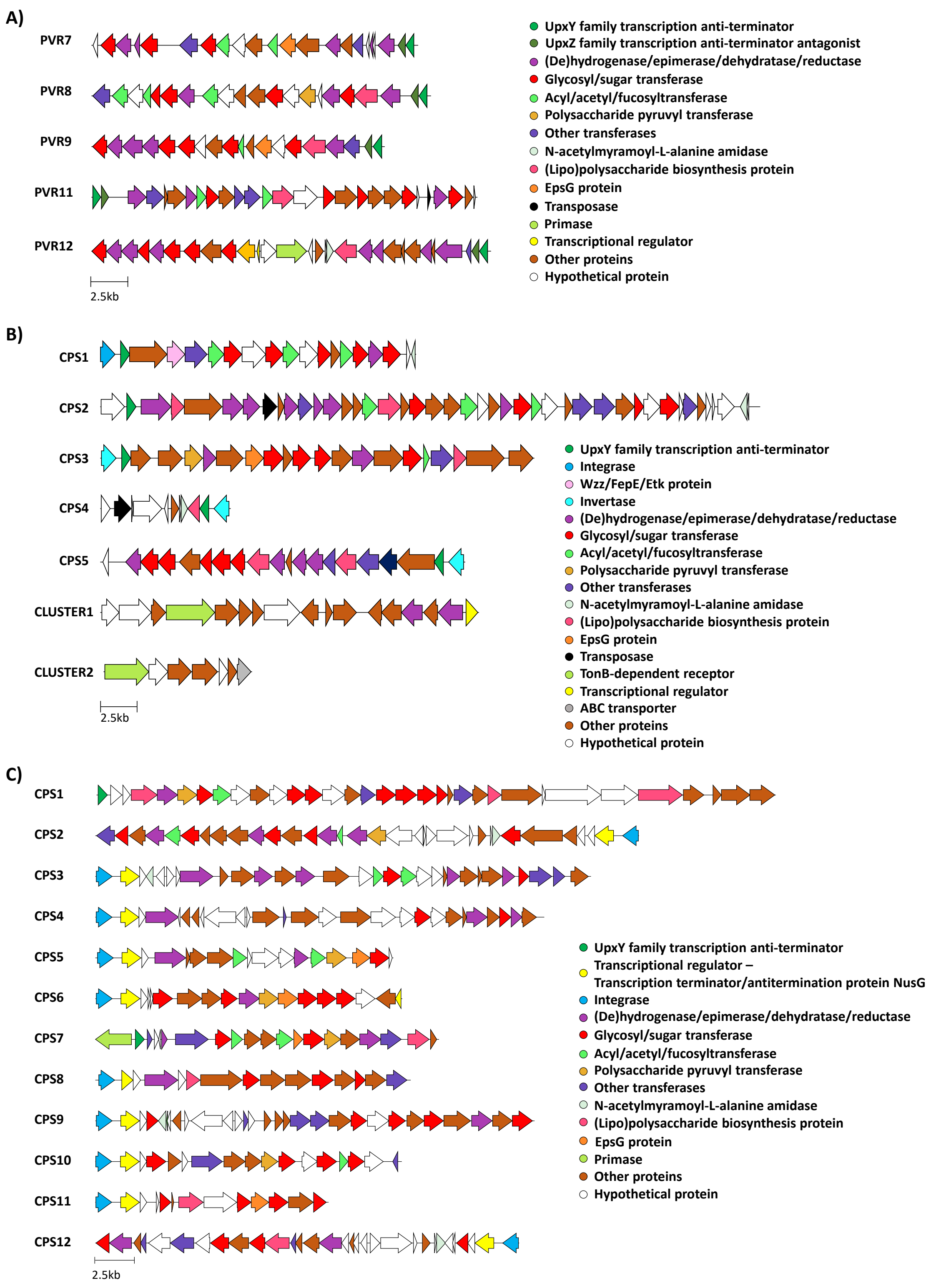


**Fig. S3.** Representation of the capsule polysaccharide (CPS) operons and gene clusters, which showed significant changes in expression after phage infection. **(A)** *B. intestinalis*, **(B)** *B. xylanisolvens* and **(C)** *P. distasonis*.

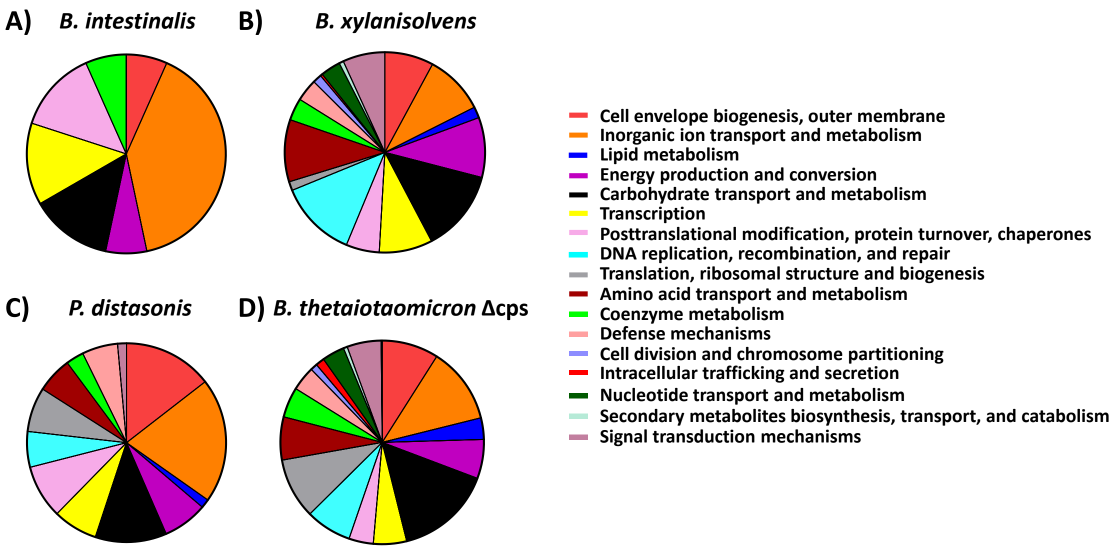


**Fig. S4.** Functional classification of the proteins encoded by differentially expressed genes identified in RNA-seq analysis using the Cluster of Orthologous Groups (COGs) database. The classification and categorisation of protein functions represented are based on genes with expression changes that were located outside of CPS loci. Each graph represents the percentage distribution of protein functions according to COG categories with known functions. **(A)** *B. intestinalis*, **(B)** *B. xylanisolvens*, **(C)** *P. distasonis*, **(D)** *B. thetaiotaomicron* Δcps.

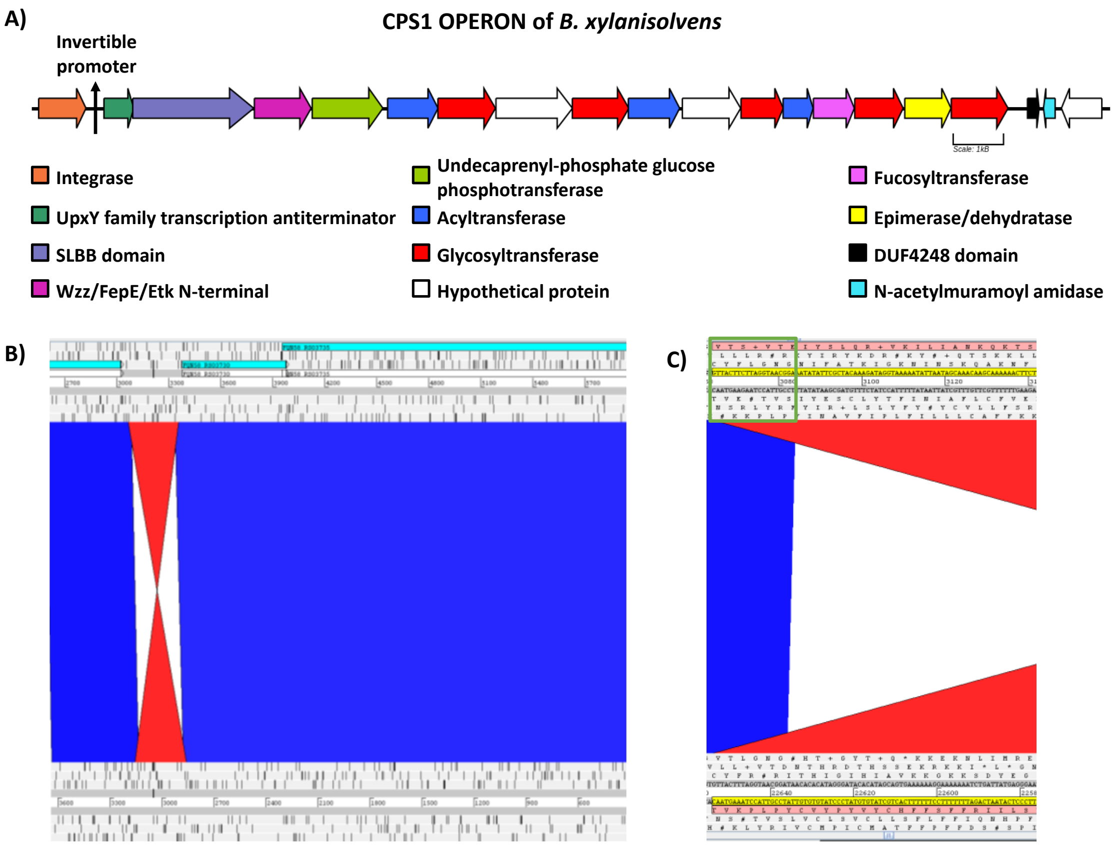


**Fig. S5.** Phase-variable region and structural inversion of CPS1 in *B. xylanisolvens*. **(A)** Structure of the phase-variable operon encoding CPS1 in *B. xylanisolvens*, with each gene represented as an arrow coloured according to the gene product, as indicated in the legend. The location of the invertible promoter is also shown. **(B)** The intergenic region of the CPS1 operon shows an inversion when long-read sequencing data are compared to the reference genome. The blue area represents regions of perfect alignment, while the red area indicates inversely aligned sequences. **(C)** Close-up view of the intergenic region where the inversion occurs, displaying the nucleotide sequence involved in this structural variation (GTTACTTCTTAGGTAACGGA).

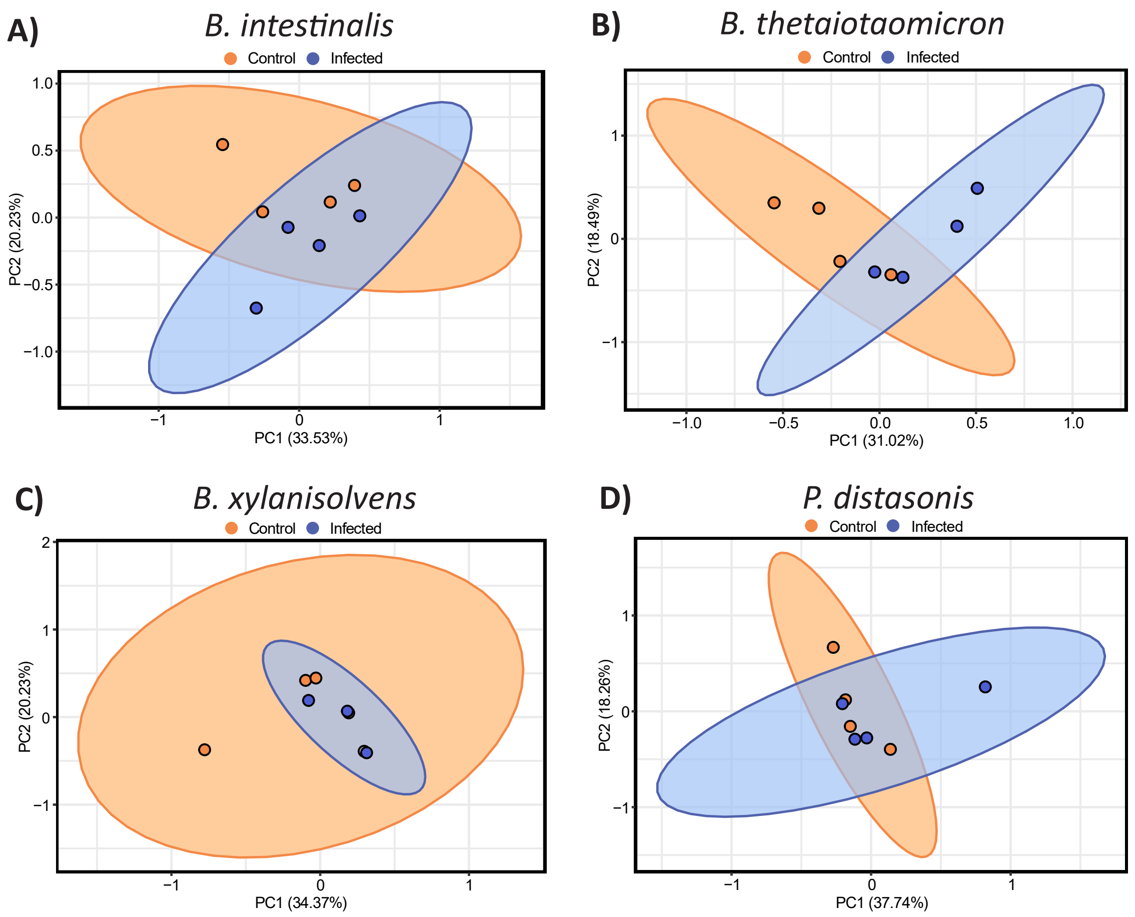


**Fig. S6.** Principal Component Analysis (PCA) of proteomic profiles obtained from the pellet during the mid-log phase on the fifth day of the co-culture experiment. The orange colour represents the control condition (uninfected), while blue is the infected condition. Ellipses represent 95% confidence intervals, and the explained variances of the two principal components are shown in brackets along the axes. **(A)** *B. intestinalis* + ϕcrAss001, **(B)** *B. thetaiotaomicron* Δcps + DAC15, **(C)** *B. xylanisolvens* + ϕcrAss002, **(D)** *P. distasonis* + ϕPDS1.

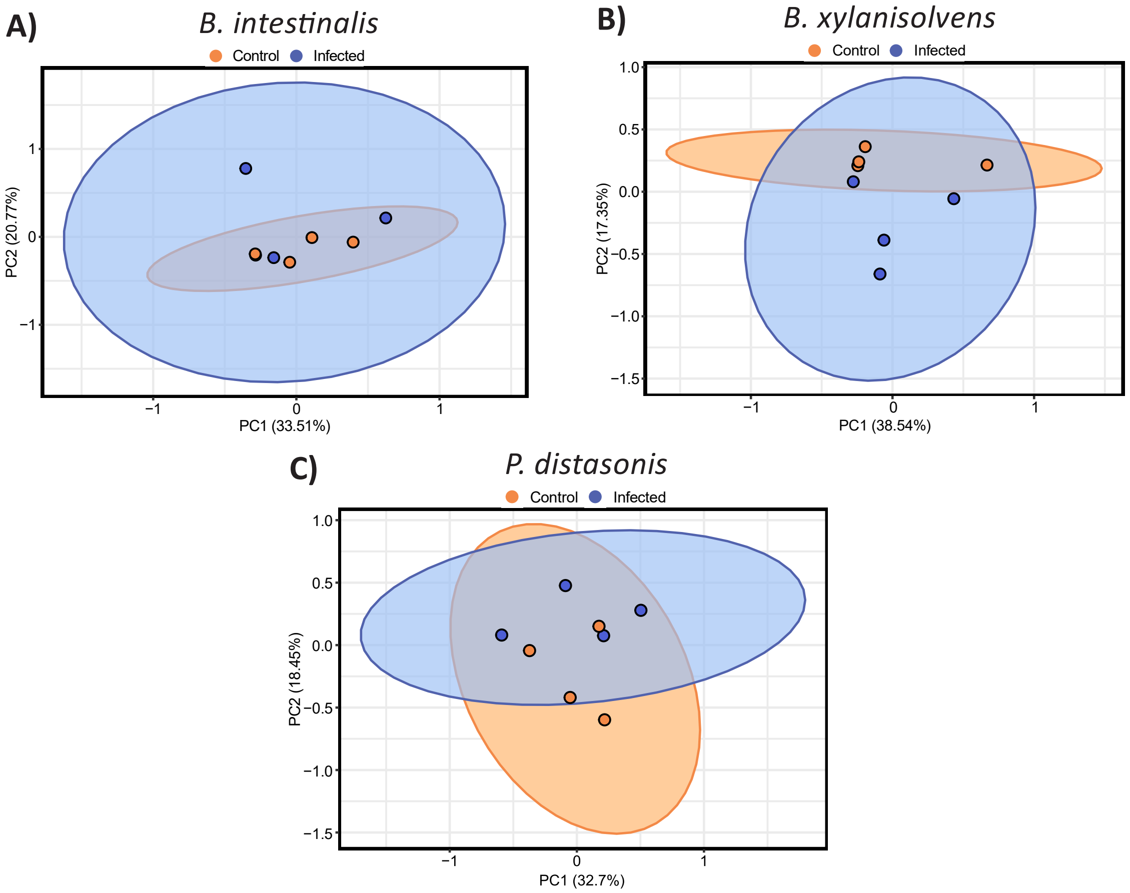


**Fig. S7.** Principal Component Analysis (PCA) of proteomic profiles obtained from the supernatant during the mid-log phase on the fifth day of the co-culture experiment in capsular bacterial strains. Orange colour represents the control condition (uninfected), while blue represents the infected condition. Ellipses represent 95% confidence intervals, and the explained variances of the two principal components are shown in brackets along the axes. **(A)** *B. intestinalis* + ϕcrAss001, **(B)** *B. xylanisolvens* + ϕcrAss002, **(C)** *P. distasonis* + ϕPDS1.

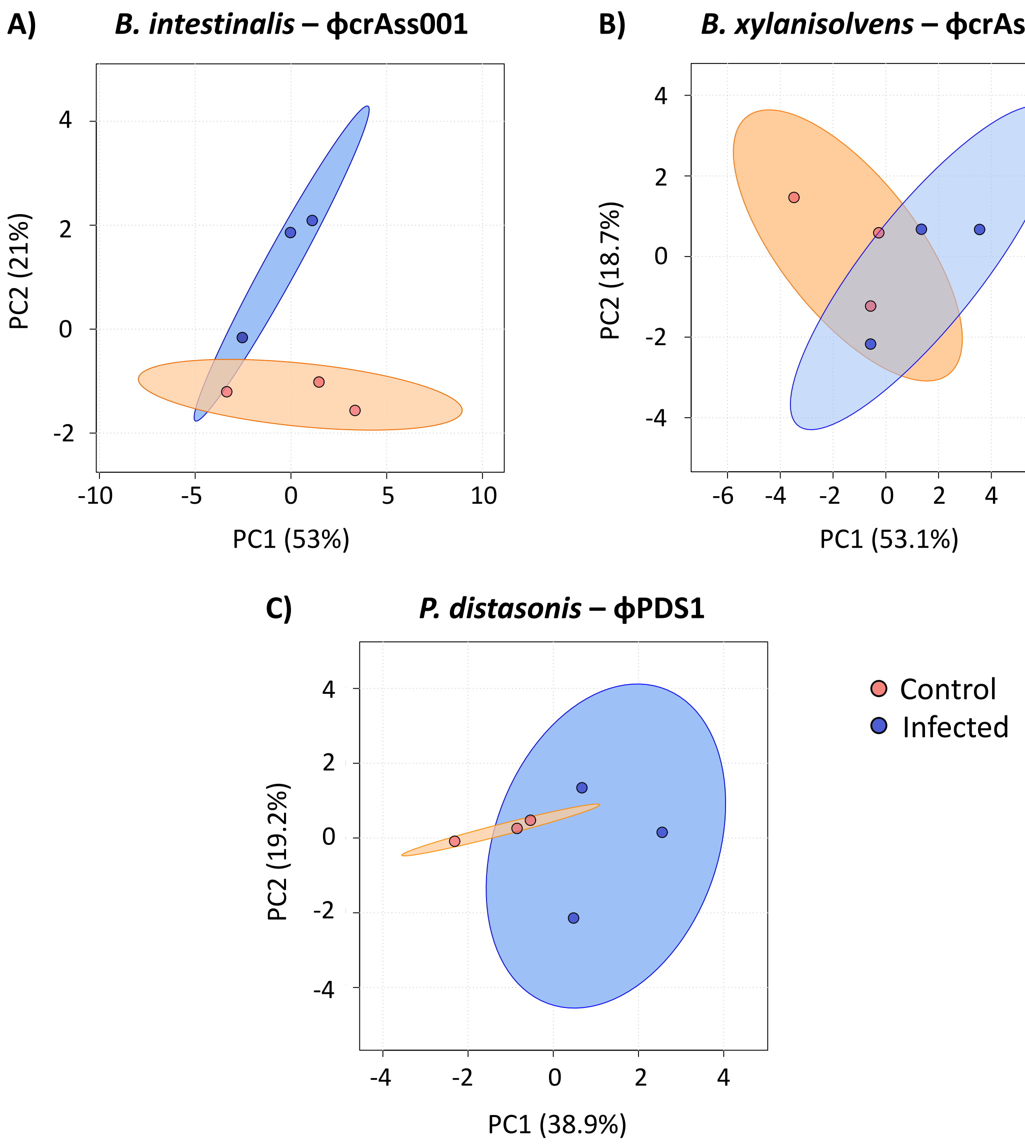


**Fig. S8.** Principal Component Analysis (PCA) of intracellular metabolomic profiles of detected compounds with accuracy levels 1 and 2a, extracted from the bacterial pellets at the mid-log phase on the fifth day of the co-culture experiment. Orange represents the control condition (uninfected), while blue is the infected condition. Ellipses represent 95% confidence intervals, and the explained variances of the two principal components are shown in brackets along the axes. **(A)** *B. intestinalis* + ϕcrAss001, **(B)** *B. xylanisolvens* + ϕcrAss002, **(C)** *P. distasonis* + ϕPDS1.
